# An engineered culture vessel and flow system to improve the *in vitro* analysis of volatile organic compounds

**DOI:** 10.1101/2023.08.05.552027

**Authors:** Jarrett Eshima, Taylor R. Pennington, Youssef Abdellatif, Angela Ponce Olea, Joel F. Lusk, Benjamin D. Ambrose, Ethan Marschall, Christopher Miranda, Paula Phan, Christina Aridi, Barbara S. Smith

## Abstract

Volatile organic compounds (VOCs) are a biologically important subset of an organism’s metabolome, yet *in vitro* techniques for the analysis of these small molecules vary substantially in practice, restricting the interpretation and reproducibility of study findings. Here, we present an engineered culture tool, termed the “Biodome”, designed to enhance analyte sensitivity by integrating dynamic headspace sampling methodology for the recovery of VOCs from viable biological cultures. We validate the functionality of the device for *in vitro* volatile metabolomics utilizing computational modeling and fluorescent imaging of mammalian cell culture. We then leverage comprehensive two-dimensional gas chromatography coupled with a time-of-flight mass spectrometer and the enhanced sampling capabilities afforded by our tool to identify seven VOCs not found in the media or exogenously derived from the sampling method (typical pitfalls with *in vitro* volatilome analysis). We further work to validate the endogenous production of these VOCs using two independent approaches: (i) glycolysis-mediated stable isotopic labeling techniques using ^13^C_6_–D-glucose and (ii) RNA interference (RNAi) to selectively knockdown β-oxidation via silencing of *CPT2*. Isotope labeling reveals 2-Decen-1-ol as endogenously derived with glucose as a carbon source and, through RNAi, we find evidence supporting endogenous production of 2-ethyl-1-hexene, dodecyl acrylate, tridecanoic acid methyl ester and a low abundance alkene (C17) with molecular backbones likely derived from fatty acid degradation. To demonstrate applicability beyond mammalian cell culture, we assess the production of VOCs throughout the log and stationary phases of growth in ampicillin-resistant DH5α *Escherichia coli*. We identified nine compounds with results supporting endogenous production, six of which were not previously associated with *E. coli*. Our findings emphasize the improved capabilities of the Biodome for *in vitro* volatile metabolomics and provide a platform for the standardization of methodology.

## Main

Volatile metabolites, part of the broad class of compounds known as volatile organic compounds (VOCs), are products of cellular metabolism with low molecular weight and a sufficient vapor pressure to exist as a gas at standard temperature and pressure. VOCs have long been recognized to confer information about biological processes in health and disease^1–3^, yet reliable methods for the study of these molecules *in vitro* have lagged behind. Works overcoming the challenges associated with *in vitro* sampling methodology have provided evidence for the extensive roles of volatile metabolites in biology, including but not limited to, serving as putative disease biomarkers^4–6^, extracellular signaling molecules in archaea, fungi, bacteria, protists, plants, and animals^7–16^, and drivers of inter– and intra–kingdom phenotype and function^7, 9–11, 17–20^. Despite the insights gained, nearly all approaches suffer from one or more of the following major limitations: i) detection of exogenous VOCs from plastics or the sampling environment^18, 21, 22^, ii) low analyte sensitivity and poor signal-to-noise relative to VOCs originating from non-endogenous sources, as a consequence of indirect^18, 23^ and passive (equilibrium-based) sampling techniques^21, 24, 25^ and/or iii) disrupt culture viability^26–28^, limiting interpretation in response to environmental stimuli or perturbation. Thus, *in vitro* VOC studies would largely benefit from a standardized method and tool to reduce signal from contaminants, enhance sensitivity, and maintain viability for time-dependent applications.

Previous groups have recognized these limitations and worked to develop a broad family of approaches known as purge-and-trap or dynamic headspace sampling (DHS)^4, 29–33^. These approaches involve the continuous sampling or removal of the headspace, often trapping the VOCs on sorbent material fixed to solid support^33–36^, or directly analyzing the extracted gas by coupling the sampling methodology to a mass spectrometer^37–39^ or sensor array^24, 40^. DHS has been shown to yield a significant increase in signal when considering the volatilome emitted from fruit extract^41, 42^ and other complex matrices^43^. Despite the advantages of and advances in DHS methodology, translation to an *in vitro* biological setting has been limited and a standardized approach does not yet exist^4, 18, 29–31, 44^. Challenges contributing to this gap include organism viability and material compatibility, exogenous contaminants, imaging capabilities, liquid-to-headspace ratio and signal sensitivity, gas flow characteristics and reproducibility, reusability, throughput, and adaptability for automation.

Here, we worked to address many of these gaps surrounding the analysis of VOCs from *in vitro* cultures through the design, development, and application of a custom borosilicate glass culture vessel (termed the “Biodome”) and demonstrate the advantages of our tool for biological analysis through the unbiased characterization of mammalian cell and bacterial volatilomes. Similar to a recently published approach^32^, this work leverages the enhanced sensitivity, resolution, and separation capabilities of thermal desorption coupled to comprehensive two-dimensional gas chromatography with time–of–flight mass spectrometry (GC×GC-TOFMS) for metabolomic analysis^45^. We pair this instrumentation with an offline, open flow system designed to trap culture-derived VOCs using a tri-phase thermal desorption tube. The modular aspects of our system support “offline” culturing in a standard incubator and easy replacement of the input gas for adaptation to oxygen-sensitive organisms. The borosilicate glass culture vessel enables sterilization by autoclaving while also allowing for imaging directly through the device.

Given the intended biological applications, we first use fluorescent imaging to demonstrate that SK-OV-3 mammalian cell growth is equivalent to a standard T-75 flask. We then work to extensively characterize the exogenous VOCs originating from the flow system and culture media in order to identify seven VOCs strictly observed in the SK-OV-3 ovarian adenocarcinoma *in vitro* volatilome, four of which have not been previously reported. Leveraging glycolysis-mediated isotopic labeling strategies^46^, we help verify 2-Decen-1-ol as cellularly derived with glucose as a carbon source. Both *in vitro* and *in vivo* mammalian volatilomes typically show hydrocarbons as a major constituent. Therefore, to validate endogenous origin using an independent approach, we apply RNA interference (RNAi) to selectively knockdown lipid metabolism through silencing of *CPT2* and identify at least four VOCs with evidence supporting biosynthesis derived from lipid metabolism. Finally, we extend the functionality of our tool by characterizing the *Escherichia coli (E. coli*) volatilome across the log and stationary phases of growth and identify six low abundance VOCs not previously observed in *E. coli* volatilome, with established pathways putatively giving rise to their biosynthetic origin. Through this work, we demonstrate that our tool enables the reproducible sampling of complex biological matrices and provides sufficient sensitivity to allow for the identification of endogenous volatile metabolites and low abundance VOCs not present in the culture media or flow system.

### Design and Development

The biological VOC sampling system is comprised of a circular glass culture vessel, compressed gas cylinder and dual stage regulator, Nalgene^TM^ 890 FEP tubing (Catalog # 8050-0310, Thermo Scientific, Waltham, MA, USA), oxygen-compatible PTFE tape (Catalog # 22485, Restek, Bellefonte, PA, USA), hydrocarbon trap (Catalog # 22013, Restek), flow meter (Catalog # 40402-0010, Riteflow, Bel-Art, Wayne, NJ, USA), sterile filter (0.2 µm pore size, Pall Corporation, Port Washington, NY, USA), bead bath (LabArmor, Plano, TX, USA), Carbopack C, Carbopack B, and Carbosieve SIII sorbent thermal desorption tube (TDT; Gerstel, Linthicum, MD, USA), and inexpensive laboratory-made TDT adapter – incorporating a modular design to facilitate adoption for a variety of biological organisms (**Fig. 1a – d**). For example, the bead bath temperature can be raised to promote growth of thermophiles or replaced with an ice bath to allow study of psychrophiles. Similarly, the inflow gas can be replaced to support the analysis of oxygen-sensitive microorganisms. The system is designed to enable the continuous sampling of the headspace VOCs in the Biodome by flowing the desired compressed gas through the culture vessel, effectively increasing the cumulative VOC signal as compared to static equilibrium-based systems (i.e. solid phase microextraction (SPME) in a culture vessel or closed vial), as a consequence of Henry’s Law.

**Fig. 1.**
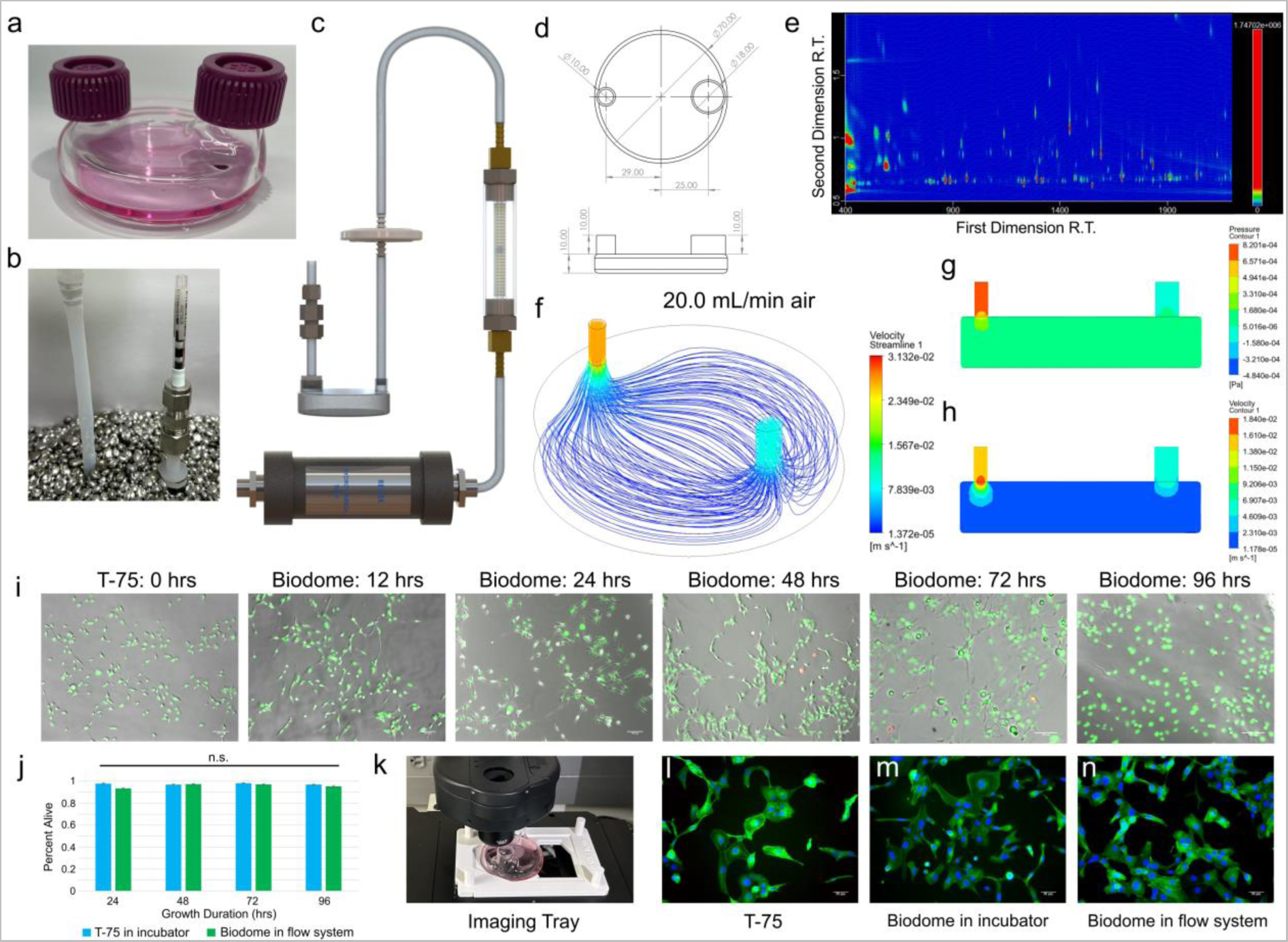
Design and development of the Biodome. (a) The borosilicate glass culture vessel, termed the Biodome, containing 5mL RPMI 1640 media. T-25 gas permeable flask caps were allowed to rest over the inlet and outlet of the vessel while in an incubator. (b) Placement of the Biodome during VOC sampling shows the glass vessel is fully submerged in beads at 37°C. Also shown is the interface between the Biodome, TDT adapter, and TDT. A sterile rubber stopper and PTFE tape are used to create an airtight seal at the interface between the TDT and the adapter. (c) 3D rendering showing the major components of the gas flow system including the hydrocarbon trap, flow meter, sterile filter, tubing, Biodome, TDT adapter, and TDT sampling tube. The compressed gas cylinder is not shown. (d) Schematic showing relevant dimensions (in mm; OD) for the Biodome culture vessel. (e) Representative GC×GC chromatogram spanning 400-2200 seconds in the first dimension and 0.5-2 seconds in the second dimension, following 24 hours of sampling SK-OV-3 ovarian adenocarcinoma cells. Fluid modeling in ANSYS^TM^ demonstrates (f) laminar flow conditions are maintained at flow rates ≤ 20 mL/min (TDT adapter insert included) and (g) gauge pressure and (h) velocity changes due to the inflowing gas are negligible at roughly half the Biodome height, limiting turbulent interactions with the surface of the cell culture media. (i) Live/dead assay images in the Biodome show viability across 0.5 – 4 days. A T-75 flask at t = 0 (24 hours post seeding) is shown for reference. (j) Independent live-dead assays were performed using NucBlue^TM^ (Hoechst 33342) and NucGreen^TM^ ReadyProbes^TM^ at 24-hour intervals, with live percentages estimated from 10 independent regions of the T-75 or Biodome culture vessels. No statistically significant differences in cell viability are observed demonstrating equivalence as compared to standard culture vessels. (k) Custom imaging tray was designed in SolidWorks^®^ 2019 and 3D printed to allow imaging using a Leica DMi8. (l-n) Typical SK-OV-3 morphology was visualized using fluorescent imaging and a live actin tracking stain (green color), Hoechst 33342 (blue color; live stain), and NucGreen (red color; dead stain), and no differences are observed using different culture vessels and growth environments. All brightfield and fluorescent images were taken using cells at passage < 10.

The Biodome culture vessel is glass blown from borosilicate glass and features a 6.35 mm OD (1/4 inch) inlet and 18mm OD outlet (**Fig. 1a, 1d**), with heights equal to 10 mm. The internal chamber height also measures 10 mm, selected to enhance VOC sensitivity and reproducibility by optimizing surface area to volume ratio while limiting turbulent interactions between the inflowing gas and the surface of the liquid culture media. The thickness of the glass is roughly 1-2 mm and the diameter of the biodome is 70 mm, giving an approximate surface area of 38.5 cm^2^. Vented caps from standard T-25 flasks were found to maintain a favorable environment for growth when allowed to rest over the inlet and outlet (**Fig. 1a**). Worth noting, no surface coating was required to promote cell adhesion and growth, increasing throughput, and reducing user burden.

The TDT adapter (**Fig. 1b, 1c, Fig. S1**) is comprised of a size 3 silicon rubber stopper adjusted to a ∼15mm diameter using a sterile razor (VWR, Radnor, PA, USA) with 6 mm hole introduced by a biopsy punch (Integra LifeSciences, Princeton, NJ, USA), Nalgene tubing (6.35 mm ID, 7.9375 mm OD, see above), one 6.35mm (1/4 inch) and one 7.9375 mm (5/16 inch) 316 stainless steel compression nut with metal ferrule (Swagelok Southwest Co., Phoenix, AZ, USA), and a 316 stainless steel 6.35 mm male to 7.9375 mm male adapter (Swagelok). For assembly of the adapter, see Methods. Although the initial design was intended for TDT sampling by two dimensional GC×GC-TOFMS (**Fig. 1e**), given the popularity of SPME for VOC analysis, we further modified the adapter in an inexpensive manner to enable this method by adding a 1 mL disposable syringe body (Electron Microscopy Sciences, Hatfield, PA), with the plunger removed, in line with the 6.35 mm (1/4 inch) nut (replacing the TDT, **Fig. S1**) and completely sealing the interface with oxygen-compatible PTFE.

To assess the gas flow characteristics through the headspace of the culture vessel, fluid modeling was performed in ANSYS Fluent^®^ 2019 R3. Results show that laminar flow is maintained for gas flow rates ≤ 20 mL/min, using the defined geometry (**Fig. 1d, 1f – h, Supplemental Video 1**). A flow rate of 11.7 mL/min was maintained for all testing, corresponding to a reading of 30 on the flow meter, unless otherwise noted. To enhance throughput, a multiplexed part was designed in SOLIDWORKS^®^ 2019 and flow characteristics were assessed using ANSYS Fluent (**Fig. S1**). Results show the part provides consistent flow rates at each of the three outlets, equal to 1/3 the total flow rate, while maintaining a single inlet (**Fig. S1, Supplemental Video 2**). Given an internal chamber volume of approximately 200 cm^3^, the internal chamber can be flushed of environmental gas using a purge flow rate of 20 mL/min and a duration of 10 minutes.

Cell viability was considered in SK-OV-3 ovarian adenocarcinoma cells (HTB-77, ATCC, Manassas, VA) using live/dead fluorescence assays. 0.5x10^6^ cells were seeded into the Biodome containing a total of 5mL of RPMI1640 culture media containing 10% FBS and 1% Pen/Strep and given 24 hours to allow cell attachment. Brightfield images show time-dependent increases in Biodome confluency and high viability (**Fig. 1i**). A standard T-75 flask (VWR) 24 hours after seeding is shown for reference, equivalent to t = 0 in the Biodome (**Fig. 1i**). Results from independent live/dead fluorescence assays (NucBlue and NucGreen ReadyProbes, Invitrogen, Carlsbad, CA) show no significant differences in SK-OV-3 cell viability in the Biodome, as compared to a standard T-75 flask in a humidified incubator, for a minimum duration of 4 days (**Fig. 1j**). We further considered cell viability through morphological fluorescence staining using a live actin tracking stain (CellMask, Invitrogen). To enable imaging on a Leica DMi8 microscrope, we designed a custom imaging tray in SOLIDWORKS^®^ 2019 (**Fig. 1k**). Following seeding in the appropriate culture vessel, SK-OV-3 cells were allowed to proliferate for 48 hours before staining, and our results show that cell morphology in the Biodome is comparable to a T-75 flask grown in a standard incubator (**Fig. 1l – n**). Taken together, our findings demonstrate that the Biodome tool maintains favorable growth conditions for at least 4 continuous days, without the need for surface treatment.

### Characterizing device performance in mammalian culture

SK-OV-3 cells (human origin) were seeded into the Biodome glass culture vessel and allowed to adhere and proliferate for 24 hours. Prior to sampling, culture media was replaced to ensure the accumulation of VOCs were minimized. Based on the previously observed growth characteristics (**Fig. 1h-1l**), cells were subject to VOC collection for 4 continuous days with the TDT replaced every 24 hours, although shorter sampling times were also shown to be viable (**Fig. S2**). RPMI 1640 was maintained as the primary medium, however dialyzed FBS was implemented in place of FBS, supported by a previous work that demonstrated serum-free growth media is favorable for VOC analysis^47^.

Results show the Biodome system supports the identification of cellular VOCs absent in the flow system and culture vessel, both with and without culture media. To focus on the reproducible aspects of the cellular and control volatilomes, we removed features observed in fewer than 3 of the 4 days of sampling, resulting in a total of 384 unique peaks (**Fig. 2a**). The filtered volatilome was subsequently subject to principal component analysis (PCA) and distinct clusters corresponding to the control and SK-OV-3 conditions were observed (**Fig. 2b**).

**Fig. 2.**
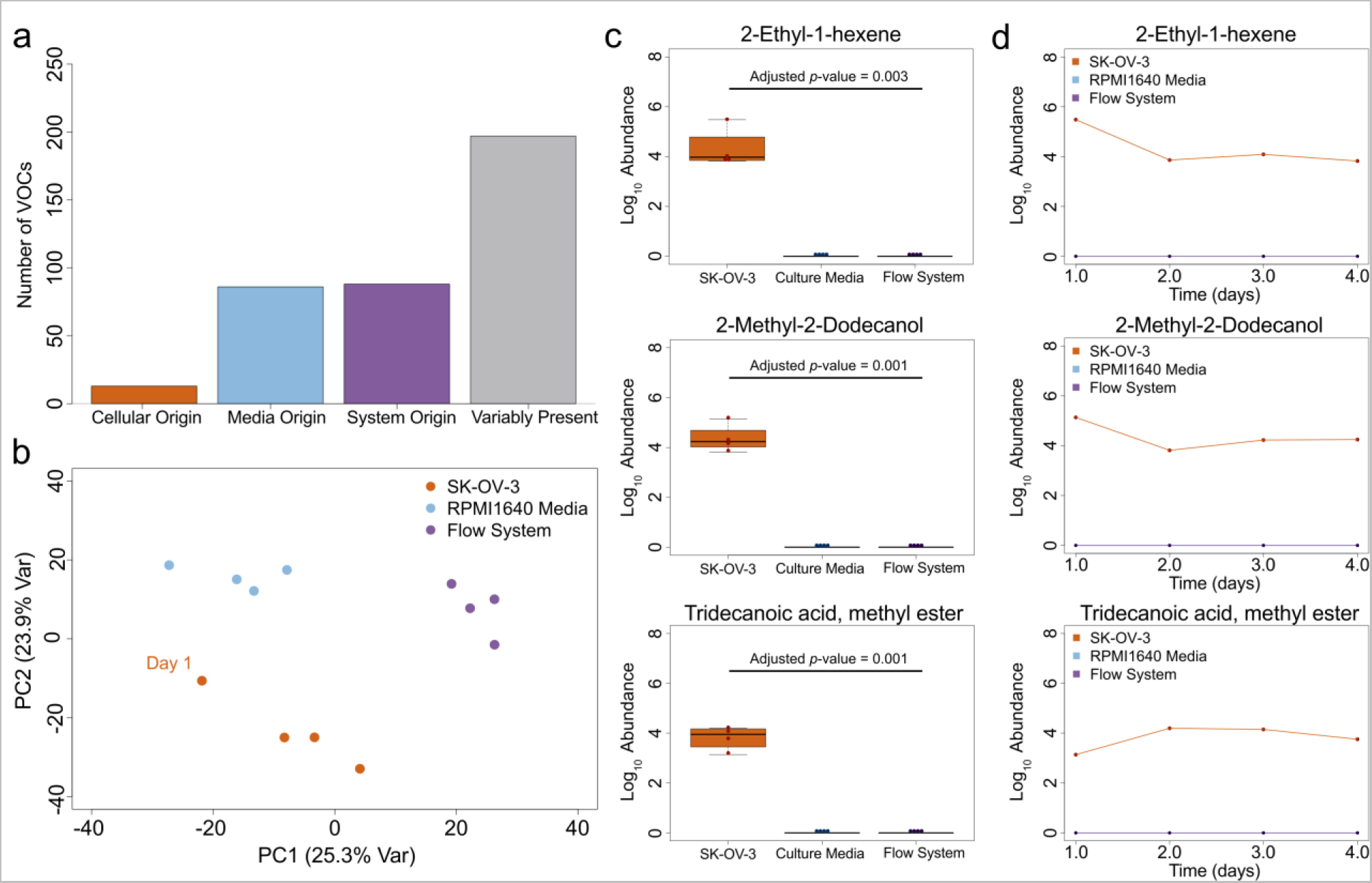
Volatilome characterization using the *in vitro* Biodome sampling tool. (a) Bar chart showing the number of VOCs detected, categorized according to their proposed origin. Variably present indicates the VOCs were observed inconsistently in two or more experimental conditions. (b) PCA plot shows distinct clusters when considering the first two principal components, using the 384 features comprising the filtered volatilome. The first day SK-OV-3 volatilome was seen to resemble the media control more closely, as compared to the other three days, suggesting a 24 “flushing” period prior to collection may help with the reproducibility of *in vitro* mammalian volatilomes. Representative (c) boxplots and (d) line plots show three low-abundance VOCs (ID < 4) reproducibly and exclusively detected in the SK-OV-3 ovarian adenocarcinoma *in vitro* volatilome. Abundance values were calculated by integrating the peak area using the unique mass and log_10_ transformed. The adjusted *p* value was calculated using a two-tailed student’s t-test and corrected using the Benjamini-Hochberg procedure^53^.

Roughly 22% (86/384) of the unique features were observed in all samples (allowing for one missing observation), including the empty Biodome control, suggesting these are inherent contaminants associated with the use of our *in vitro* tool (**Fig. 2a**, **Supplemental Dataset 1**). We further find that five VOCs capture 9–44% of the total chromatographic signal in the empty culture vessel, after removing poorly resolved peaks with a first-dimension retention time < 400 seconds (see Methods). These high abundance contaminant VOCs (naming confidence ID level^48^ ≤ 3, see Methods) were found to include: Ethyl acetate, Pyridine, 1-Methyl-2-pyrrolidinone, 3,7-Dimethyl-1-octene, and 1-Dodecanol. An alkane standard (**Fig. S3**) was run to allow identification of VOC retention indices (RI) and used to support name assignment where possible (see Methods). A further 13% (50/384) of the unique features were observed in both the culture media and SK-OV-3 experimental conditions but not the empty Biodome control, suggesting media origin (**Fig. 2a**, **Supplemental Dataset 2**). Interestingly, 38 VOCs (9.9%) were reliably detected in the media volatilome but not the SK-OV-3 volatilome, which may indicate SK-OV-3 metabolic biotransformation but more likely signal suppression, given 73.7% had a median signal-to-noise ratio < 150:1 (**Supplemental Dataset 2**). Lastly, a total of 13 VOCs (3.4%) were strictly observed in the experimental condition containing SK-OV-3 cells, allowing for 1 total spurious peak from controls. The remaining features (n=197) that passed our filtering criteria were *primarily* present in: (i) system control (122; 61.9%), (ii) media control (39; 19.8%), or (ii) SK-OV-3 condition (36; 18.3%) yet were variably present in the other two experimental groups, limiting further interpretation. Four of these VOCs were strictly observed in cellular volatilomes *and*, to the best of the authors’ knowledge, have not previously detected in mammalian cancer cells (bolded names), emphasizing the enhanced sensitivity afforded by the *in vitro* Biodome tool (**Table 1**; **Fig. 2c, 2d; Fig. S4**).

**Table 1.**
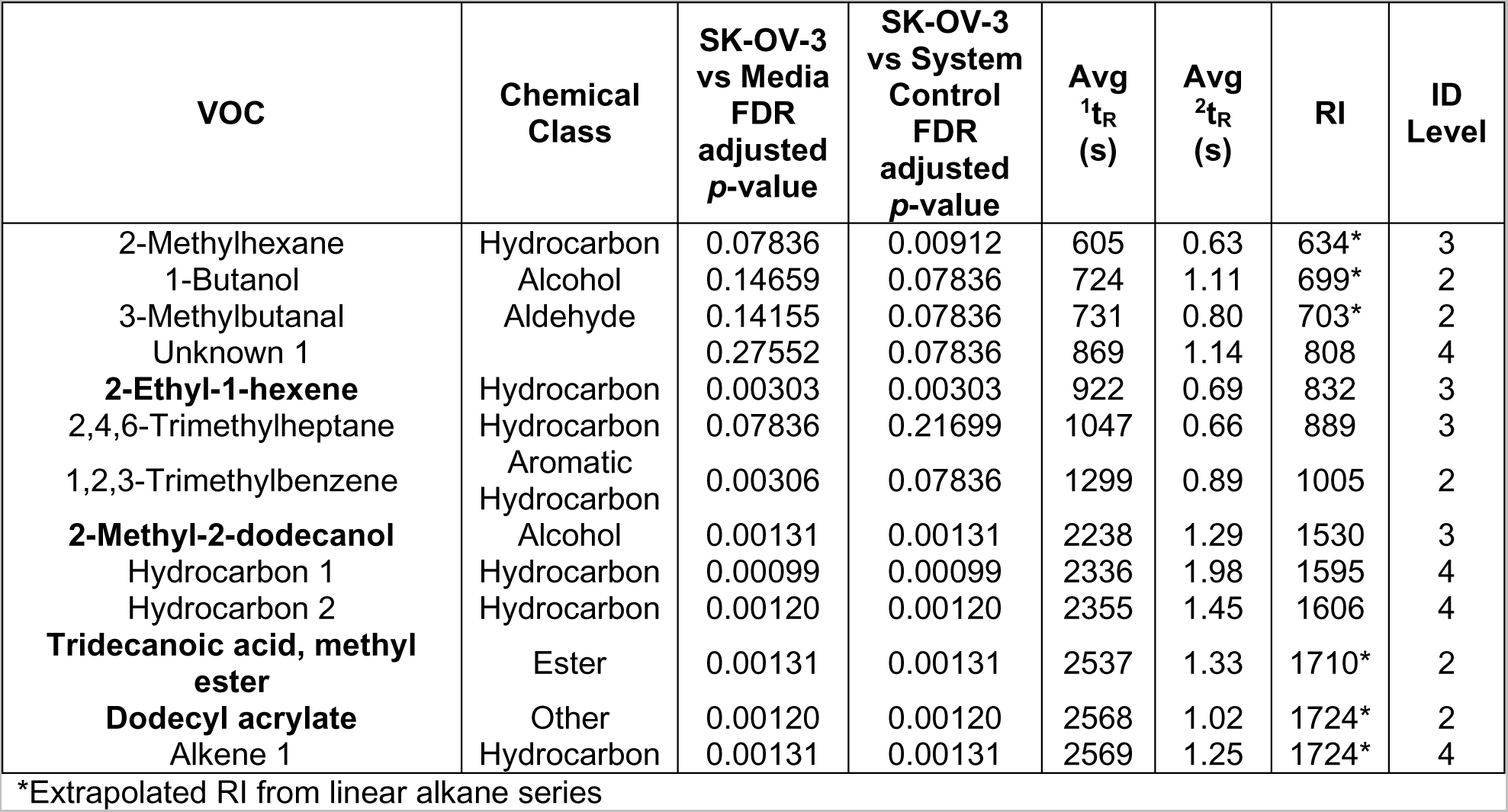
VOCs reproducibly detected from SK-OV-3 cells. The compound name is presented first, followed by the functional group class. Bolded names indicate that the VOC was not detected in the media or system controls and have not been previously reported *in vitro*. The adjusted *p* value was calculated using a two-tailed student’s t-test, comparing the control and SK-OV-3 volatilomes post-filtering. The first (^1^t_R_) and second (^2^t_R_) dimension retention times are included, with the retention index calculated using a C8-C20 alkane standard mix. Finally, the VOC naming confidence level is reported according to previously established standards.

Cell-specific VOCs were primarily hydrocarbons but spanned multiple chemical classes, including the reproducible detection of three VOCs that fell below the naming threshold. Despite the lower naming confidence, mass spectral data could be used to support their identification as two alkanes eluting at 2336 and 2355 seconds, and an alkene eluting at 2569 seconds, in the first dimension (**Table 1**, **Fig. S5**). Importantly, while mass spectra indicate alkane (*m/z* 43, 57, 71, 85, etc.) and alkene (*m/z* 41, 55, 69, 83, etc.) backbones, second dimension retention times suggest they may be functionalized. Of the VOCs observed strictly in the SK-OV-3 volatilomes, over half (54%) had a median signal-to-noise ratio < 150:1, emphasizing the capability of our approach to recover low abundance volatiles that may be missed using equilibrium-based sampling approaches.

### Stable isotope labeling of metabolites in cell culture (SILMC)

For the global and unbiased assessment of endogenous VOC production^46^, SK-OV-3 cells were passaged 20 times supplemented with ^13^C_6_–D-glucose (Cambridge Isotope Laboratories, Tewksbury, MA, USA) to isotopically label glucose-derived metabolites (**Fig. 3a**). As described previously, labeled (^13^C_6_–D-glucose) and unlabeled (^12^C_6_-D-glucose) cells were seeded into the Biodome device and allowed to grow out for 24 hours.

**Fig. 3.**
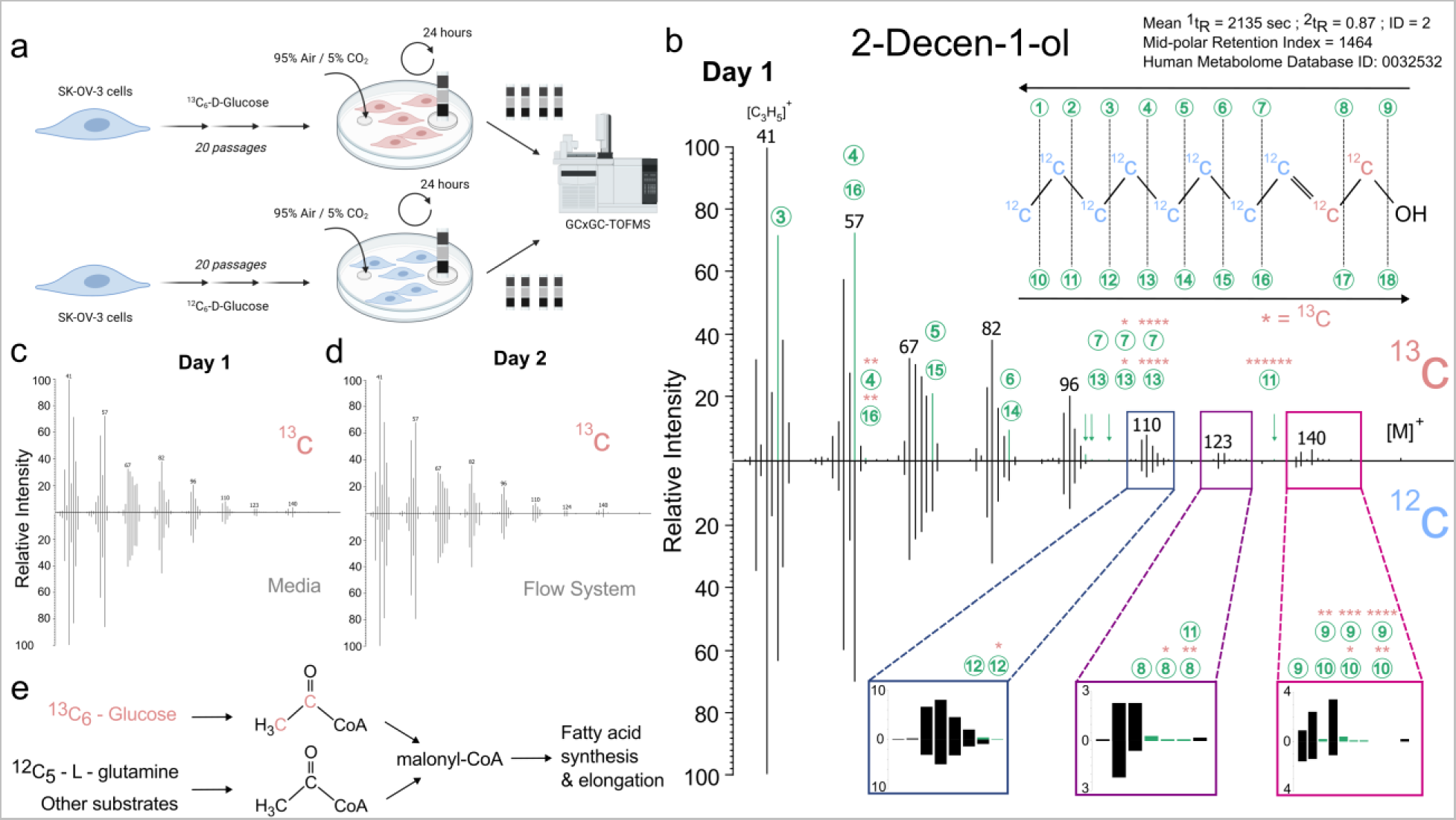
Glycolysis-mediated isotopic labeling in the Biodome. (a) Diagram showing the workflow implemented for the stable isotope labeling of glycolysis-derived VOCs. Representative mass spectrums comparing (b) labeled and unlabeled, (c) ^13^C-labeled and media control, and (d) ^13^C-labeled and flow system control were selected using the peak with the highest signal-to-noise ratio across the four days of sampling. Despite observing 2-Decen-1-ol in non-cellular control volatilomes, many low intensity fragments remained uniquely observed in the ^13^C-labeled spectra. Breakout images showing *m/z* ranges between 107 – 114, 122 – 128, and 137 – 147 amu were generated in R using raw mass spectral data. The proposed ^13^C-labeled structure is indicated in red, while annotations correspond to the unlabeled (^12^C) fragments. Fragments without an annotation are possibly derived from secondary bond rearrangements. (e) Proposed metabolic pathway for ^13^C incorporation into 2-Decen-1-ol.

Labeled and unlabeled SKOV-3 lines, at equivalent passage number, were subject to VOC collection for 4 continuous days, with TDU tubes replaced every 24 hours (**Fig. 3a**). TDU tubes were stored for a maximum of 8 days at 4°C to limit degradation^49^ prior to analysis by GC×GC-TOFMS. Chromatograms from the same experimental condition were aligned using ChromaTOF^®^, and subsequently processed using custom R code to match isotopically labeled peaks to unlabeled ones using thresholding of the first- and second-dimension retention times – given ^13^C-labeled VOCs should elute from the column at the same time as ^12^C VOCs^50, 51^. Following post-processing in R, putative ^13^C-labeled VOCs were manually validated by comparing ions and parent fragments between the ^12^C and ^13^C experimental conditions.

Our results capture one VOC, 2-Decen-1-ol, with mass spectral information supporting ^13^C-labeling derived from ^13^C_6_–glucose (**Fig. 3b**; **Table 2; Fig. S6**). We provide possible structures for mass fragments primarily or uniquely detected in the ^13^C-labeled spectra (**Table 2**). Further evidence supporting our claims include, (i) chromatographic elution of labeled and unlabeled 2-Decen-1-ol was consistent, with a relative standard deviation <1% in the first- and second-dimension, (ii) the mid-polar retention index falls within the expected range for 2-Decen-1-ol, leading to a naming confidence level of 2, and (iii) 2-Decen-1-ol has been previously observed to act as an endogenous metabolite in humans and observed extracellularly^52^. Despite observing 2-Decen-1-ol in media and flow system control volatilomes, many tentative ^13^C fragments remained uniquely observed in the ^13^C-labeled SK-OV-3 spectra (**Fig 3c, 3d; Fig. S6**). Analyte fragments of *m/z* 103, 107, 114, 125, 126, 127, 133, 141, 142, 143 and 147 were strictly observed in the ^13^C-labeled spectra, corresponding to the addition of 2 carbon isotopes near the alcohol functional group, with spectral and metabolic evidence supporting carbon positions 1 and 2 (**Fig 3e**).

**Table 2.** Putative ^13^C fragments of 2-Decen-1-ol. Isotope fragments are presented first increasing in mass, followed by the relative intensity in the unlabeled (^12^C) and ^13^C-labeled 2-Decen-1-ol mass spectra for all four days of sampling. Chemical structures potentially giving rise to the observed signal are proposed where possible, although secondary rearrangements cannot be ruled out. Further, mass fragments without a listed structure are reasoned to be derived from secondary bond rearrangements.

| Isotope Fragment (amu) | Relative Intensity in $^{12}\text{C}$-labeled (%) | Relative Intensity in $^{13}\text{C}$-labeled (%) | Possible Structure(s) |
| --- | --- | --- | --- |
| 59 | 0.7, 1.1, 0.0, 0.0 | 0.5, 0.2, 0.4, 0.5 | $^{12}\text{C}_1^{13}\text{C}_2\text{H}_5\text{O}$ |
| 91 | 0.6, 0.0, 0.8, 0.0 | 0.3, 0.2, 0.2, 0.2 |  |
| 100 | 0.0, 0.0, 0.0, 0.0 | 0.1, 0.0, 0.0, 0.2 | $^{12}\text{C}_6^{13}\text{C}_1\text{H}_{15}$ ; $^{12}\text{C}_5^{13}\text{C}_1\text{H}_{11}\text{O}$ |
| 103 | 0.0, 0.0, 0.0, 0.0 | 0.2, 0.0, 0.0, 0.0 | $^{12}\text{C}_2^{13}\text{C}_4\text{H}_{15}$ ; $^{12}\text{C}_2^{13}\text{C}_4\text{H}_{11}\text{O}$ |
| 107 | 0.0, 0.0, 0.0, 0.0 | 0.1, 0.0, 0.2, 0.0 |  |
| 114 | 0.0, 0.0, 0.0, 0.0 | 0.1, 0.0, 0.0, 0.0 | $^{12}\text{C}_6^{13}\text{C}_1\text{H}_{11}\text{O}$ |
| 125 | 0.0, 0.0, 0.0, 0.0 | 0.3, 0.0, 0.2, 0.3 | $^{12}\text{C}_9\text{H}_{16}$ |
| 126 | 0.0, 0.0, 0.0, 0.0 | 0.1, 0.0, 0.0, 0.0 | $^{12}\text{C}_8^{13}\text{C}_1\text{H}_{16}$ |
| 127 | 0.0, 0.0, 0.0, 0.0 | 0.1, 0.0, 0.0, 0.0 | $^{12}\text{C}_7^{13}\text{C}_2\text{H}_{16}$ |
| 133 | 0.0, 0.0, 0.0, 0.0 | 0.3, 0.0, 0.0, 0.0 | $^{12}\text{C}_2^{13}\text{C}_6\text{H}_{15}\text{O}$ |
| 139 | 0.0, 1.3, 0.0, 0.0 | 0.2, 0.4, 0.0, 0.2 | $^{12}\text{C}_{10}\text{H}_{19}$ |
| 141 | 0.0, 0.0, 0.0, 0.0 | 0.4, 0.0, 0.0, 0.5 | $^{12}\text{C}_8^{13}\text{C}_2\text{H}_{19}$ ; $^{12}\text{C}_9\text{H}_{17}\text{O}$ |
| 142 | 0.0, 0.0, 0.0, 0.0 | 0.1, 0.3, 0.0, 0.0 | $^{12}\text{C}_7^{13}\text{C}_3\text{H}_{19}$ ; $^{12}\text{C}_8^{13}\text{C}_1\text{H}_{17}\text{O}$ |
| 143 | 0.0, 0.0, 0.0, 0.0 | 0.1, 0.0, 0.0, 0.0 | $^{12}\text{C}_6^{13}\text{C}_4\text{H}_{19}$ ; $^{12}\text{C}_7^{13}\text{C}_2\text{H}_{17}\text{O}$ |
| 147 | 0.0, 0.0, 0.0, 0.0 | 0.2, 0.0, 0.0, 0.0 |  |

**Table 3.** *E. Coli* and LB Broth VOCs with an ID level < 4. The compound name is presented first, followed by the functional group class. Bolded names indicate that the VOC was not detected in the broth controls. The adjusted *p* value was calculated using a two-tailed student’s t-test, comparing the LB broth and *Escherichia coli* volatilomes post-filtering. The first (^1^t_R_) and second (^2^t_R_) dimension retention times are included, with the retention index calculated using a C8-C20 alkane standard mix. Finally, the VOC naming confidence level is reported according to previously established standards.

| VOC | Chemical Class | FDR<br>adjusted<br>*p*-value | Avg<br><sup>1</sup>t<sub>R</sub><br>(s) | Avg<br><sup>2</sup>t<sub>R</sub><br>(s) | RI | ID<br>Level |
| --- | --- | --- | --- | --- | --- | --- |
| Acetone | Ketone | 0.5939 | 639 | 0.67 | 652* | 2 |
| **Methacrolein** | Aldehyde | 0.0420 | 702 | 0.69 | 687* | 2 |
| **1-Propanol** | Alcohol | 0.0278 | 709 | 1.02 | 691* | 2 |
| **Cyclobutane** | Hydrocarbon | 0.0769 | 799 | 1.06 | 758* | 3 |
| 2,3,5-Trimethylhexane | Hydrocarbon | 0.6285 | 994 | 0.65 | 865 | 3 |
| 2,3-Hexanedione | Ketone | 0.4370 | 1003 | 0.85 | 869 | 2 |
| 2,3,4-Trimethylhexane | Hydrocarbon | 0.6684 | 1028 | 0.66 | 880 | 3 |
| Nonane | Hydrocarbon | 0.6285 | 1084 | 0.65 | 906 | 2 |
| 5-Methyl-2-hexanone | Ketone | 0.6720 | 1141 | 0.88 | 932 | 2 |
| **2-Methyl-3-hexanone** | Ketone | 0.0118 | 1156 | 0.82 | 939 | 3 |
| 2,5-Dimethylpyrazine | Heteroaromatic | 0.6500 | 1199 | 1.11 | 959 | 2 |
| Cyclohexanone | Ketone | 0.8560 | 1232 | 1.05 | 974 | 2 |
| Decane | Hydrocarbon | 0.0584 | 1326 | 0.67 | 1019 | 2 |
| Dimethyl trisulfide | Other | 0.2917 | 1358 | 1.09 | 1034 | 2 |
| Trimethylpyrazine | Heteroaromatic | 0.2161 | 1369 | 1.06 | 1040 | 2 |
| 1,2,3-Trimethylbenzene | Aromatic Hydrocarbon | 0.8560 | 1370 | 0.90 | 1040 | 2 |
| 2-Ethyl-5-methylpyrazine | Heteroaromatic | 0.5463 | 1377 | 1.03 | 1044 | 2 |
| 2,2,7,7-Tetramethyloctane | Hydrocarbon | 0.5660 | 1392 | 0.66 | 1051 | 3 |
| 2,2,4,4-Tetramethyloctane | Hydrocarbon | 0.7921 | 1395 | 0.67 | 1053 | 3 |
| 2,3,5,8-Tetramethyldecane | Hydrocarbon | 0.7679 | 1438 | 0.67 | 1073 | 3 |
| 2-Acetylthiazole | Azole | 0.1401 | 1462 | 1.53 | 1085 | 2 |
| 2-Methyl-3-isopropylpyrazine | Heteroaromatic | 0.5613 | 1482 | 0.97 | 1095 | 2 |
| **Benzyl alcohol** | Functionalized Benzene | 0.0133 | 1510 | 0.35 | 1109 | 2 |
| 3-Ethyl-2,5-dimethylpyrazine | Heteroaromatic | 0.3556 | 1526 | 0.99 | 1118 | 2 |
| 2-Hydroxybenzaldehyde | Other | 0.5586 | 1534 | 1.54 | 1122 | 2 |
| 1-Octanol | Alcohol | 0.8560 | 1536 | 1.11 | 1123 | 2 |
| **2-Nonanone** | Ketone | 0.0094 | 1581 | 0.86 | 1146 | 2 |
| Benzoic acid, methyl ester | Functionalized Benzene | 0.9380 | 1605 | 1.23 | 1159 | 2 |
| 2-Methylundecane | Hydrocarbon | 0.5178 | 1628 | 0.70 | 1171 | 2 |
| 3,5-Diethyl-2-methylpyrazine | Heteroaromatic | 0.4370 | 1666 | 0.94 | 1191 | 2 |
| (E)-2-Nonenal | Aldehyde | 0.5242 | 1729 | 0.99 | 1225 | 2 |
| 1-Phenyl-1-propanone | Functionalized Benzene | 0.5178 | 1760 | 1.24 | 1242 | 2 |
| Benzyl nitrile | Functionalized Benzene | 0.5713 | 1766 | 1.76 | 1246 | 2 |
| 3-Butyl-2,5-dimethylpyrazine | Heteroaromatic | 0.5178 | 1848 | 0.94 | 1291 | 2 |
| **1-Phenyl-2-butanone** | Functionalized Benzene | 0.0088 | 1881 | 1.21 | 1310 | 2 |
| Indole | Heteroaromatic | 0.0097 | 2100 | 1.20 | 1442 | 2 |
| 3-hydroxy-2,2,4-trimethylpentyl<br>isobutyrate | Other | 0.1074 | 2129 | 1.00 | 1460 | 3 |

\*Extrapolated RI from linear alkane series

Importantly, our mass spectral data indicates variable ^13^C labeling, which may be a consequence of the inherently low ^13^C-labeled fraction and/or multi-step biosynthesis routes, with detectable mass shifts between 1-6 amu depending on the sampling day (**Table 2**). Furthermore, a ^13^C-shifted parent ion beyond an *m/z* of 156 was not observed, however given the parent peak had a median relative intensity of 0.5% across all spectra, it is likely that any labeled fraction fell below the detectable limit. To enhance detectable isotope fraction, future studies may benefit from using multiple ^13^C-labeled carbon sources (e.g. L-glutamine)^52^.

### Gene knockdown supports endogenous production of cellular VOCs

To highlight the viability and analyte sensitivity benefits of *in vitro* volatolomics in the Biodome, we investigated the relationship between the biosynthetic origin of cellular VOCs and *CPT2* transcriptional levels in SK-OV-3 cells. The *CPT2* expression levels were altered by treating the SK-OV-3 cells with pooled siRNA oligonucleotides mediating gene knockdown (see Methods). The expression levels of the *CPT2* relative to the housekeeping gene *GAPDH* was determined using RT-qPCR. Results show *CPT2* expression was significantly reduced in siRNA-treated SK-OV-3 cells to 24.3 ± 7.01% (mean ± S.E.M; p < 0.05) at t = 0 hrs, indicating successful transfection and gene knockdown (**Fig. 4a**). The expression levels of *CPT2* in the transfected SKOV-3 cells were also monitored in a 6-well plate format over a 96-hour time period. As depicted in **Fig. 4b**, the *CPT2* expression level during the initial 24 hours showed a significant decrease to 9.83 ± 4.20% (p < 0.05). Subsequently, after 48 hours, the expression levels increased to 53.0 ± 7.82% (p < 0.05). Following 72 hours, the expression levels decreased to 36.8 ± 6.49% (p < 0.05) and were recorded as 56.8 ± 10.5% (p < 0.05) after 96 hours.

**Fig. 4.**
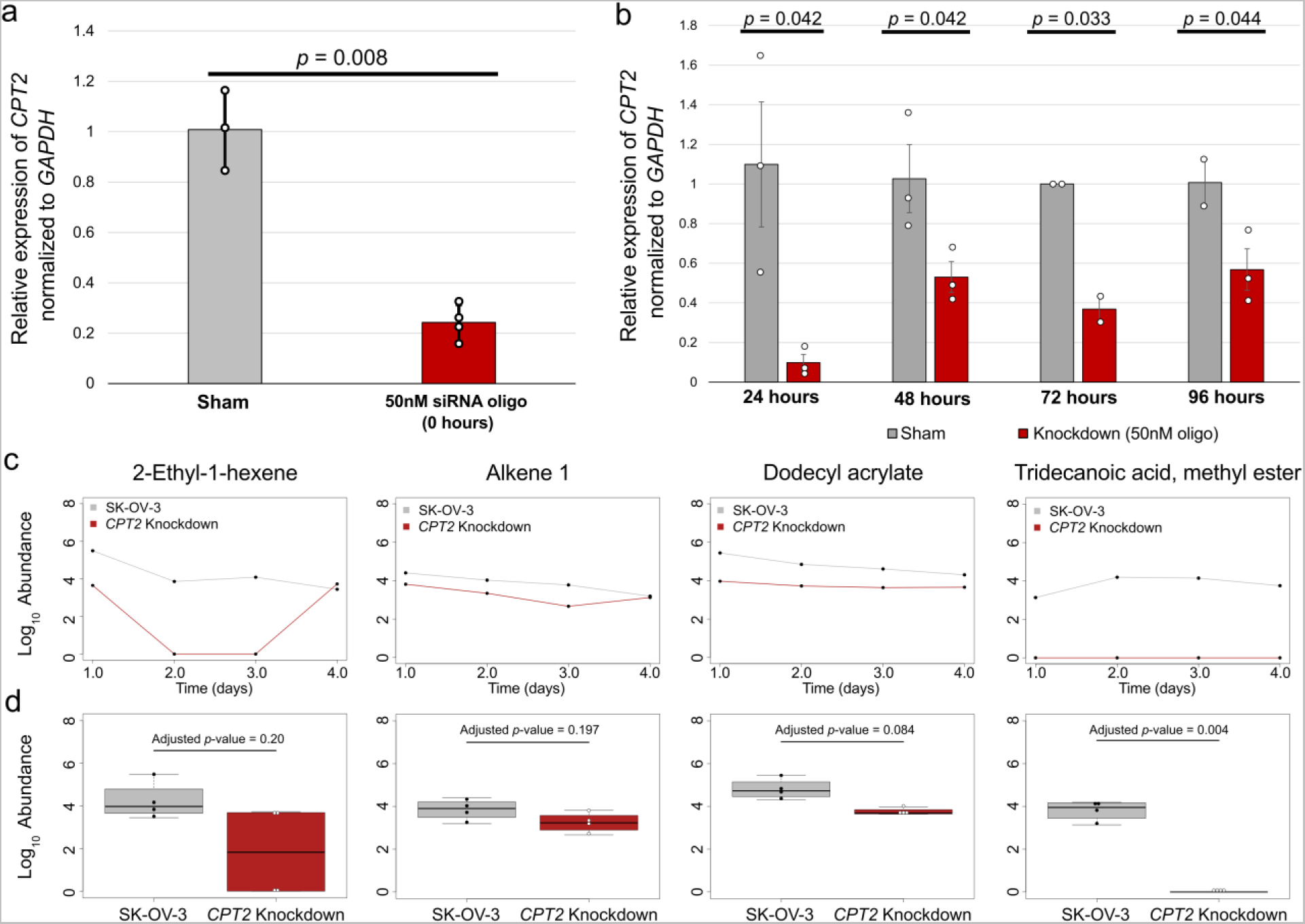
Knockdown of *CPT2* supports endogenous VOC production. (a) Expression of *CPT2* quantified by RT-qPCR at the start of VOC collection (time = 0) using biologically independent replicates. (b) The expression of *CPT2* was independently mapped across the four days of VOC sampling following siRNA-mediated knockdown in SK-OV-3 cells at equal passage number using biologically independent replicates. (c) Four VOCs not found in controls showing decreased abundance is consistent with the transient inhibition of β-oxidation via *CPT2* knockdown, across 4 days of measurement. (d) Boxplots showing significant or trending toward significant decreases in endogenous VOC abundance following integration by unique mass and log_10_ transformation. *P*-values were adjusted using the Benjamini-Hochberg procedure^53^.

Associated with the knockdown of *CPT2* in SK-OV-3 transfected cells, at least four VOCs were found to have decreased abundances consistent with the transient inhibition across the 96-hour time period (**Fig. 4c**, 4d**; Fig. S7; Supplemental Dataset 3**). In the case of 2-Ethyl-1-hexene, the median decrease in raw peak abundance was found to be 77%, with a roughly 98.5% decrease observed during the first day of sampling (**Fig. 4c, 4d**). Similarly, the unknown alkene abundance was observed to have a median decrease of 78.8%, with a 74% reduction on the first day, followed by 79%, 92%, and 15% on days 2, 3, and 4 respectively. Dodecyl acrylate showed a 91.1% decrease in the median abundance following knockdown of *CPT2* with reductions of 97%, 92%, 89%, and 77% at days 1-4, respectively. Observations at each time point are consistent with the transient recovery of *CPT2* following siRNA-mediated knockdown suggesting this VOC is directly linked to lipid β-oxidation (**Fig. 4b, 4c**). Although tridecanoic acid, methyl ester was not observed in the *CPT2* knockdown volatilome, this VOC was determined to have low abundance with a median signal-to-noise ratio of 30 across the four days of sampling. Our data processing method set a signal-to-noise threshold at 5 (see Methods), therefore the greatest reduction in signal while still allowing detection would be ∼85%, suggesting that the lack of observation may still support association with lipid metabolism given the inherently low abundance. Although signal-to-noise thresholding may partially explain the difference in reported abundance, we observe a significant decrease in tridecanoic acid, methyl ester (adjusted *p* = 0.004) and a trend towards significance (adjusted *p* = 0.084) in the case of dodecyl acrylate after correction for multiple hypothesis testing. Interestingly, production of dodecyl acrylate and the alkene were consistent in SK-OV-3 cells despite knockdown of *CPT2*, with the highest amount of signal detected on the first day, suggesting these VOCs may be associated with cellular growth and proliferation.

### Bacterial adaptation

To extend the functionality of our tool, we analyzed VOCs originating from *Escherichia coli,* strain DH5α, across 3 days, during the log and stationary phases of growth. *E. coli* were transformed with plasmid containing ampicillin resistance gene and seeded into the Biodome containing LB Broth and 100 µg/mL ampicillin. Turbid broth, indicating an increase in cell density, was observed at the conclusion of the sampling period for the run containing *E. coli* but not in the broth control (**Fig. S8**). Following *in vitro* VOC sampling and analysis by GC×GC-TOFMS, chromatographic peaks were aligned in ChromaTOF^®^, yielding 196 unique features after the removal of known contaminants and compounds observed in the instrument blanks. For statistical analysis, compounds not detected in at least 80% of all samples were removed to focus on the reproducible aspects of the *E. coli* volatilome (see Methods, **Supplemental Dataset 4**). VOCs specific to the *E. coli* or broth volatilome were also retained, resulting in a total of 108 features (**Table S1**). Results show the Biodome has sufficient sensitivity to identify low abundance bacterial-derived VOCs, even when initially present in the culture media (**Fig. 5**, **Fig. S8**).

**Fig. 5.**
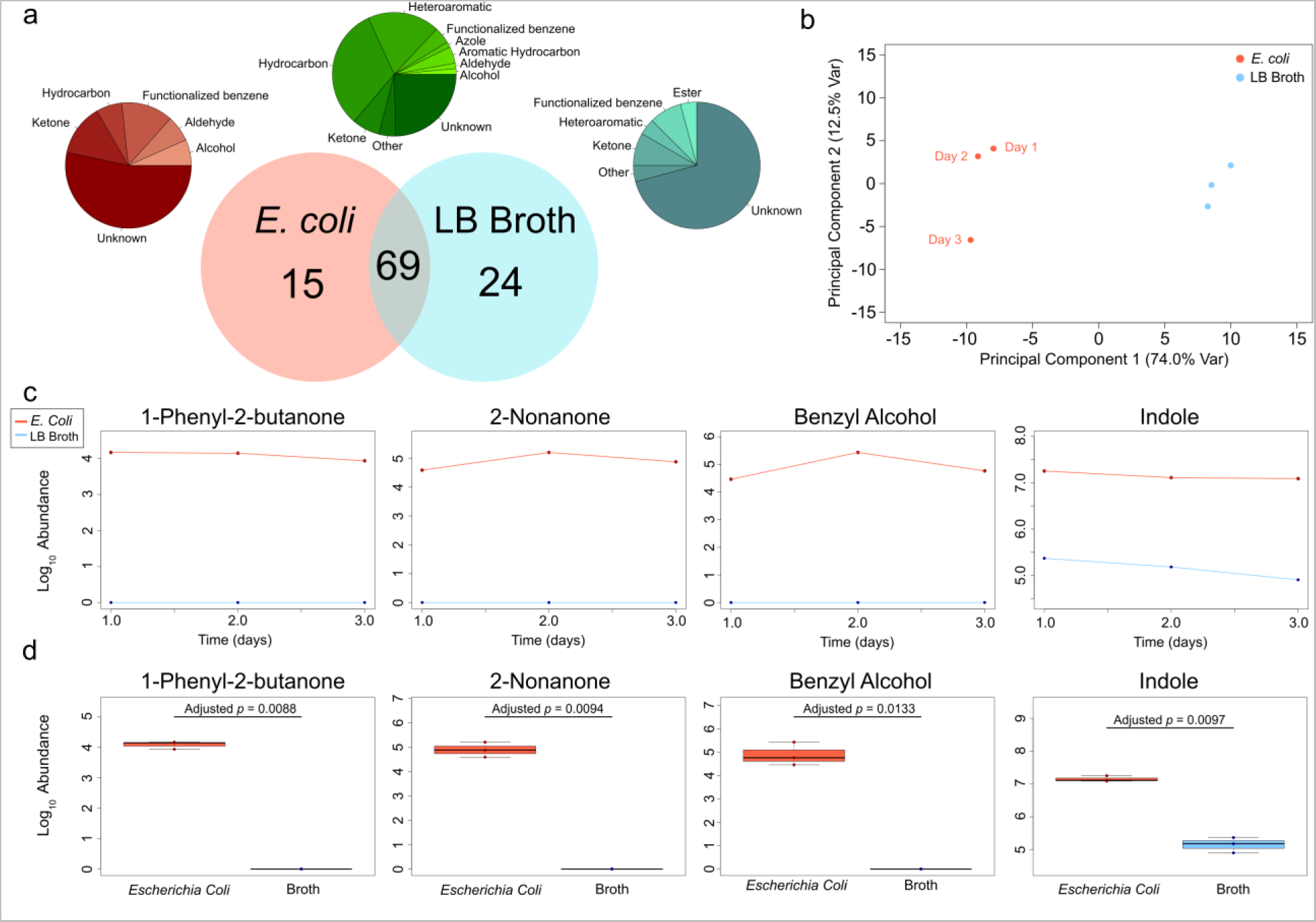
Application of the Biodome tool to characterize the *E. coli* volatilome across the log and stationary phases of growth. (a) Venn diagram shows the number of unique and common VOCs detected in the bacterial and broth conditions. Pie charts show the breakdown of VOC functional groups specific to each group. (b) PCA analysis using high confidence VOCs presented in Table 2 shows the *E. coli* volatilome is distinct from the broth control. (c) Line plots for four high confidence VOCs showing log_10_ transformed abundance across three days of sampling. Day one represents the cumulative volatilome between 0-24 hours of growth, day 2 between 24-48 hours, and the final day capturing 48-72 hours. (d) Boxplots for the same four VOCs showing expression of these metabolites are significantly elevated in the *E. coli* experimental condition, after correcting for multiple hypothesis testing using the Benjamini-Hochberg *p*-value adjustment procedure^53^.

Using the reporting standards^48^ mentioned previously (see Methods), filtered VOCs were assigned a name and functional group where possible (**Fig. 5a**). Higher confidence VOCs with an ID level < 4 are provided in **Table 2**. VOCs uniquely observed in the broth may suggest consumption by the bacteria during log and stationary phases of growth, although additional work is needed to validate endogenous origin (**Fig. 5a**). Using the filtered bacterial and broth volatilomes, we performed PCA and observe highly separated groups when considering the first two components (**Fig. 5b**). Interestingly, spatial arrangement shows that the *E. coli* volatilome produced during the first 48 hours of growth is relatively consistent and suggests the third day of sampling may capture the transition to the death phase (**Fig. 5b**). After adjusting for multiple hypothesis testing using the Benjamini-Hochberg procedure^53^, 16 VOCs were found to be significantly elevated in the condition containing ampicillin-resistant *E. coli* (**Fig. 5c, 5d**; **Fig. S8**; **Table 2**; **Table S1**).

The majority of the identified volatiles were found in both the broth and bacterial conditions (63.9%), however abundance changes were observed that indicate not all shared VOCs originate from exogenous sources. Lending strength to claims of enhanced sensitivity, indole (FDR *p*-value = 0.010) and decane (FDR *p*-value = 0.058) were elevated in the samples containing *E. coli* despite also being observed in the liquid broth volatilome, supporting endogenous production (**Fig. 5c, 5d**). Furthermore, noise due to plastics (xylenes and styrene) accounted for less than 1% of the total chromatographic signal in each replicate, emphasizing that the Biodome effectively addresses one of the major existing limitations with existing approaches (**Supplemental Dataset 4**). Finally, of the 15 VOCs found uniquely in the *E. coli* volatilome, roughly 75% (11/15) had a median signal-to-noise ratio < 150:1, across the log and stationary phases of growth, demonstrating the analyte sensitivity advantages of our tool for untargeted volatile metabolomics.

Many of the VOCs observed in the *E. coli* volatilome have been previously linked to this microorganism, however we first report the production of low abundance VOCs 1-phenyl-2-butanone, 1-propanol, 2-methyl-3-hexanone, benzyl alcohol, cyclobutane, and methacrolein during the log and stationary phases of growth (**Fig. 5c, 5d**; **Table 2; Fig. S8**). All newly reported VOCs had a median signal-to-noise ratio < 150:1. We also briefly consider VOCs distinguishing the log phase from the stationary phase of growth and note distinct production of 1-butanol, a dimethylpyrazine isomer eluting after 2,5-dimethylpyrazine, and an unknown compound with average first- and second-dimension retention times of 940 seconds and 1.05 seconds respectively, corresponding to a retention index of 841 (**Fig. S8**).

## Discussion

In this study, we first work to design and develop a glass culture vessel to facilitate the *in vitro* analysis of biological volatilomes. We validate the functionality of our tool by integrating computational modeling and fluorescent imaging to demonstrate desirable sampling and growth conditions. We then extensively characterize the origin of VOCs from the flow system itself, the culture media, and human-derived SK-OV-3 ovarian adenocarcinoma cells. In doing so, we identify at least two compounds not previously observed *in vitro* but reported *in vivo*. To further verify endogenous origin, and demonstrate the variety of methods supported by our tool, we work to globally label ^13^C_6_– D-glucose-derived VOCs and identify 2-Decen-1-ol with mass spectral data supporting ^13^C incorporation. With a similar aim, we inhibit lipid metabolism leveraging RNAi-mediated knockdown of *CPT2* and find four additional volatile metabolites that show a decrease in abundance consistent with the transient recovery of *CPT2* expression across the 4 days of analysis. Finally, to broaden the scope of use for *in vitro* volatile metabolomics, we characterized the DH5α *Escherichia coli* volatilome across the log and stationary phases of growth (0-72 hours) and identify six low abundance VOCs not previously reported.

Considering the newly reported SK-OV-3 cell volatile metabolites^54^, 2-ethyl-1-hexene (*3-methyleneheptane*) has not been previously detected *in vitro*, however multiple studies report observing this VOC in fecal samples^55^ and in the breath of both healthy individuals^56^ and in patients following hyperbaric oxygen therapy^57, 58^, potentially linking this VOC to cellular respiration. Interestingly, following ^13^C-glucose supplementation, we observe a variable but consistent shift in the parent ion (*m/z* 112) corresponding to 1-6 ^13^C molecules incorporated (113–118 amu), however the original parent fragment is inconsistently observed limiting interpretation. Further to the contrary, 2-ethyl-1-hexene has been reported as an atmospheric VOC^59^, indicating exogenous sources cannot be ruled out entirely, although the lack of observation in media and flow system controls supports cellular origin. To the best of the authors’ knowledge, very little has been reported on 2-methyl-2-dodecanol as a mammalian metabolite^54, 56^. However 1-dodecanol, similar in structure, has been shown to act as a pheromone in non-mammalian organisms^60, 61^ – potentially indicating a relationship to the ovarian adenocarcinoma cell line utilized in this study. Tridecanoic acid methyl ester has not been previously reported in mammalian cell culture *in vitro*, however both the free fatty acid (tridecanoic acid) and esterified form have been detected from healthy human skin^56, 62^. Similarly, dodecyl (*lauryl*) acrylate has not been reported *in vitro* but has been detected from human skin by direct PDMS contact^63^. Dodecyl acrylate is known to be produced by the condensation of 1-dodecanol (HMBD0011626) with acrylic acid (HMBD0031647), both of which are known human metabolites^52^, potentially indicating its role as a secondary marker for these metabolites. To the contrary, 1-dodecanol was one of the more abundant contaminants present in the flow system, suggesting observation may strictly be an indicator of endogenous acrylic acid, although additional work is needed to verify this origin. In general, our results demonstrate that the Biodome effectively enhances total signal recovery for *in vitro* mammalian volatilomics and allows reproducible detection of low-abundance, intracellular VOCs.

Mammalian cell VOCs worth further consideration include 1,2,3-trimethylbenzene (*hemimellitene*), 1-butanol, 2,4,6-trimethylheptane, 2-methylhexane, and 3-methylbutanal (*isovaleraldehyde*). 1,2,3-trimethylbenzene has been reported *in vitro* in growth media controls and significantly elevated by *in situ* SPME and gas subsampling in A549 lung carcinoma cells^21^, suggesting endogenous origin. Here, our results show additional evidence supporting cellular origin and further find agreement in published literature with 1,2,3-trimethylbenzene reported in healthy human feces and breath^56^. Furthermore, 1,2,3-trimethylbenzne has been linked to recurrent wheezing in children and gastrointestinal disease in adults, suggesting a role in human disease^55, 64^. 1-Butanol has been previously reported *in vitro* in A549 lung carcinoma cells^65^, although some limitations to biological interpretation exist as covered by Schallschmidt et al.^21^. Our results provide additional evidence for endogenous origin and our findings are further supported by the detection of 1-butanol in most human biofluids^56^ with increases reported in the urine and serum of diabetic patients^66^, suggesting a role in human disease. To the best of the authors’ knowledge, 2,4,6-trimethylheptane has not been previously observed in the *in vitro* mammalian volatilome. However, this VOC has been reported in the breath of healthy individuals^56^, lending support to our findings. 2-Methylhexane has been previously reported *in vitro*^21^, with decreases in abundance relative to the media control, suggesting exogenous origin and/or bioconversion by the cells. Here we find evidence to the contrary – given a single observation across media and flow system controls and consistent detection in the cellular volatilome. In agreement with our findings, 2-methylhexane has been reported in the breath of healthy individuals^56^ and in patients with lung cancer, acting as a discriminatory marker for the disease^67, 68^, suggesting a role as a human metabolite. Interestingly, 3-methylbutanal has been previously observed *in vitro* from NCI-H2087 lung adenocarcinoma cells using an early dynamic headspace sampling approach and large bioreactor system developed by Sponring et al.^4^. Results from Sponring et al. indicate exogenous presence in the culture media / bioreactor, while our own findings support intracellular origin given the lack of observation in controls. The authors hypothesize differences in culture conditions, nutrient availability, and the optimization of headspace-to-volume ratio in the Biodome may partially explain the apparent differences.

The results from isotopic labeling with ^13^C_6_–D-glucose confirm 2-Decen-1-ol as an endogenous VOC derived from glycolysis. 2-Decen-1-ol is a fatty alcohol that falls under a class of molecules synthesized from elongation of an acetyl-CoA substrate. In line with established lipid biosynthetic pathways, our mass spectral data supports the incorporation of ^13^C-labeled acetyl-CoA with carbon isotopes near the alcohol functional group. Subsequent metabolic events leading to the production of 2-Decen-1-ol include chain elongation of acetyl-CoA and, eventually, fatty alcohol formation via fatty acyl-CoA reductase. Complex labeling patterns observed for 2-Decen-1ol arise from variable degrees of ^13^C incorporation in each of these biosynthetic pathways. Extensively labeled molecules corresponding to larger mass shifts depict higher degrees of ^13^C incorporation. Small mass shifts indicating a lower degree of ^13^C incorporation could potentially arise from the elongation of unlabeled fatty acids and precursors that also feed into lipid metabolism, and/or a lower degree of ^13^C-labeled acetyl-CoA added during elongation. Fluctuating degrees of ^13^C-labeled fatty acids were also observed by Kamphorst, et al., who used both ^13^C-labeled glucose and glutamine to study the kinetics of fatty acid metabolism^69^. 2-Decen-1-ol and other fatty acid metabolites are energy-rich compounds that can be utilized for cellular signaling, membrane formation, and energy production^70, 71^. Furthermore, a considerable number of studies indicate that these biological processes are altered in cancer cells to promote survival and tumorigenesis^70–72^. Notably, upregulated fatty acid metabolism plays a key role in ovarian cancer cell survival, and the expression of genes encoding acetyl-CoA metabolic enzymes is associated with increased aggressiveness and poor prognosis^71, 72^. Taken together, these findings highlight the ability of the Biodome tool to effectively capture VOCs of key biological processes *in vitro*, using an untargeted approach to identify metabolites associated with disease.

Like 2-Decen-1-ol, many VOCs detected from mammalian culture *in vitro* often feature a relatively long hydrocarbon backbone and are generally thought to originate from fatty acid metabolism. To more generally evaluate the role of fatty acid metabolism in the production of VOCs *in vitro*, we elected to transiently inhibit expression of mitochondrial lipid transporter protein carnitine palmitoyltransferase 2 (encoded by *CPT2*) using RNA interference. 2-Ethyl-1-hexene, dodecyl acrylate, tridecanoic acid methyl ester, and an unidentified alkene were strictly observed in cellular volatilomes and demonstrated abundance decreases consistent with the knockdown of *CPT2* across 4 days. As would be expected, all compounds feature a hydrocarbon backbone. 2-Ethyl-1-hexene and the unidentified alkene include a double bond which may indicate origin from an unsaturated fatty acid, while dodecyl acrylate and tridecanoic acid methyl ester are likely derived from saturated fatty acids. To the best of the authors’ knowledge, this represents the first application of RNAi to assess pathway origin of VOCs in mammalian cells, however similar success has been recently demonstrated in plants^73, 74^. Although a specific enzymatic step cannot be assigned to their production, our findings strengthen claims for cellular origin and provide a methodological platform for the secondary verification of VOC producing pathways in mammalian volatilomes.

In the case of the *E. coli* volatilome, the significant increase in indole, relative to the broth control, is an indication of the advantages of the Biodome tool for *in vitro* volatile metabolomics. Conversion of tryptophan to indole by tryptophanase is a well characterized process in *E. coli*^75, 76^. Tryptophanase catalyzes the production of pyruvate in conditions of excess tryptophan, as expected in starter broth cultures^77^, generating indole as a byproduct^75, 76^. Here, we find indole is the most abundant peak across the log and stationary phases of growth (0-72 hours) in the *E. coli* volatilome, although one recent study has shown bacterial transformation to induce ampicillin resistance may act as a confounding factor^78^. Generally, our results agree with many previous studies, which demonstrate indole is a biomarker for *E. coli*, acting as an extracellular signaling molecule with import and export complexes, and serving to regulate biofilm formation, cell division, and gene expression^79–83^. Worth discussing, indole was detected with high confidence in the broth controls and had similarly broad peaks. Therefore, to ensure consistency, peaks within the first-dimension retention time window of 2092-2110 seconds were summed in all chromatograms, likely explaining the high reported signal in broth controls despite clear differences in abundance (**Fig. S8**). Demonstrating similar agreement, recent studies also report 2-nonanone in the *E. coli* volatilome, including the DH5α strain we detected it from in this study^84, 85^. Further, biosynthesis of 2-nonanone is known to be naturally derived from the decarboxylation of 10 carbon β-keto acids^86^ and has been shown to regulate gene expression associated with quorum sensing^87^. Taken together, our findings demonstrate that volatile profiles acquired using the Biodome tool accurately reflect active metabolic processes in *E. coli* and agree with previous studies in regard to the abundance of these VOCs relative to the remainder of analytes comprising the volatilome.

While we detected the well characterized *E. coli* VOCs indole and 2-nonanone, we first report the observation of 1-phenyl-2-butanone, 2-methyl-3-hexanone, 1-propanol, benzyl alcohol, cyclobutane, and methacrolein in the DH5α *E. Coli* volatilome but not in the LB broth control. Interestingly, while 1-phenyl-2-butanone has not been previously observed *in vitro*, the structural analog 3-hydroxy-1-phenyl-2-butanone has been observed in other microorganisms^88, 89^ with a natural biosynthesis pathway reported via the condensation reaction of phenylpyruvic acid with acetaldehyde, catalyzed by phenylpyruvate decarboxylase^90^. Alternatively, oxidation of *trans*-1-phenyl-1-butene by rat cytochrome P-450 has been shown to produce 1-phenyl-2-butanone. Given previous works, the authors propose further catabolism of 3-hydroxy-1-phenyl-2-butanone to give 1-phenyl-2-butanone and/or oxidation by cytochromes – both suggesting intracellular origin. To the best of the authors’ knowledge, 2-methyl-3-hexanone has not been previously detected in the *E. coli* volatilome, although the reaction of 3-oxoacids with H^+^ to produce methyl ketones and CO_2_ is known to occur intracellularly and spontaneously^91^, supporting endogenous origin.

Further highlighting analyte recovery advantages using the Biodome, our results support the intracellular production of 1-propanol and benzyl alcohol. Most studies work to metabolically engineer *E. coli* to produce appreciable quantities of 1-propanol^92, 93^ and benzyl alcohol^94^. The former is known to be produced intracellularly as a product of β-D-glucuronide and D-glucuronate degradation^91^, isoleucine biosynthesis via 2-keto acid decarboxylase and alcohol dehydrogenase^95^, and in association with sleeping beauty mutase (Sbm) operon in wild-type *E. coli*, although expression is minimal^96, 97^. In agreement with these works, our results support intracellular production given the low abundance of 1-propanol, and further suggest detectable abundance has remained beyond the capabilities of traditional equilibrium-based sampling methodologies. Similarly, benzyl alcohol has been shown to be produced endogenously from benzaldehyde and multiple native alcohol dehydrogenases and aldo–keto reductases^94, 98^. Further, benzyl alcohol has been shown to induce heat shock proteins in *E. coli*^99^, suggesting a functional role for this volatile metabolite.

Cyclobutane has not been previously reported as a VOC originating from *E. coli*. While the natural biosynthetic routes are not well characterized^100, 101^, cyclobutane is known to be produced intracellularly during the dimerization of pyrimidine nucleotides as a consequence of UV-induced DNA damage^102^, yet repair mechanisms do not release cyclobutane^103^, potentially indicating exogenous origin. To the contrary, cyclobutane has been observed in the feces and breath of healthy humans^56^, suggesting a role as a microbial metabolite. We find additional support for intracellular origin given cyclobutane was observed with a median signal-to-noise ratio < 100:1. Finally, while little is known about the intracellular production of methacrolein, studies have shown that the non-mevalonate pathway for isoprenoid biosynthesis is active in *E. coli*^104, 105^ and that isoprene oxidation produces methacrolein^106^, with evidence supporting direct biological emission from plants^107^ and other microorganisms^108^. However, cellular origin cannot be definitively established, with evidence supporting an exogenous source including the absence of isoprene. Generally, our consideration of the *E. coli* volatilome emphasizes the analyte recovery advantages of the Biodome tool, expanding depth of coverage and enabling study of previously uncharacterized volatile metabolites.

In brief summary, our findings highlight the advantages of the Biodome tool for *in vitro* volatile metabolomic analysis, which include (i) dynamic headspace methodology to enhance total signal recovery allowing detection of lower abundance VOCs, (ii) application of borosilicate glass to allow sterilization and reuse, limit exogenous signal due to plastics, and support dual imaging function, and (iii) improved reproducibility while operating in the laminar flow range. The non-destructive nature of collection allows for analysis of VOCs across time and in response to perturbation, as demonstrated by our isotopic labeling and RNA interference applications. The modular system design facilitates adaptation to microbiological analysis, with notable usefulness for the study of anaerobes and extremophiles, and we demonstrate compatibility through the analysis of the DH5α *Escherichia coli* volatilome across the log and stationary phases of growth.

## Methods

### Biodome Development and Fabrication

A CAD model of the Biodome culture vessel, with the TDT adapter included, was designed in SOLIDWORKS^®^ 2019. Flow characteristics were then modeled in the CAD part using ANSYS Fluent^®^ 2019 R3. For meshing, the target skewness was adjusted to 0.65, smoothing set to high, a mesh metric set to “Orthogonal Quality”, and default for all other parameters. The inlet and outlet mesh were then refined using a value of 2. Boundary conditions were then set, using inlet velocities ranging from 1 – 50 mL/min and the pressure outlet condition. The operating pressure was assumed to be 1 atm, the fluid set to air, and acceleration due to gravity was included in the model. The residual threshold was adjusted to 1E-5. Solution parameters were as follows: Pressure-Velocity Coupling Scheme set to SIMPLE, Spatial Discretization Gradient set to Least Squares Cell Based, Pressure set to Second Order, and Momentum set to Second Order Upwind. Solution results were visualized using ANSYS CFD Post GUI.

Fabrication of the glass culture vessel leverages specialized trade techniques, but in brief begins with rotation of Pyrex tubing using a glassblowing lathe. A natural gas or oxygen flame was used to pull the tube down into a flat bottom. The inlet and outlet were then added to onto the flat bottom. Using a custom holder to retain that piece in the lathe, another flat bottom was pulled to create the internal chamber. Additional attention was given to the curvature of the bottom and top to ensure the flattest surface possible for imaging applications.

To assemble the TDT adapter, Nalgene tubing was inserted into the 7.9375 mm nut head (with moderate force) and the other end was inserted into the rubber stopper. A metal ferrule and tubing insert was attached to the free end and the nut-ferrule assembly was then connected to the male-to-male adapter (**Fig. S1e**). A ferrule was then added to the 6.35 mm stainless steel nut and screwed firmly into the other end of the male-to-male adapter. The opening of the 6.35mm stainless-steel nut served as the interface for TDT sampling. An airtight seal was ensured by wrapping 7-8 cm of oxygen-compatible PTFE tape around the beveled end of the TDT tube before inserting it into the TDT interface. A visible decrease in the flow rate (> 2 mL/min) was always observed when the TDT was inserted into the adapter and the regulator was readjusted to achieve the original flow rate after roughly 5-10 minutes of equilibration. To adapt the system for SPME sampling methodology, a 1 mL disposable syringe body (Electron Microscopy Sciences, Hatfield, PA), with the plunger removed, in line with the 6.35 mm (1/4 inch) nut (replacing the TDT, **Fig. S1f**) and completely sealing the interface with oxygen-compatible PTFE. The SPME fiber then rests naturally at the narrow end of the syringe body – without contacting the sides of the syringe. Other SPME adapters were considered, including the use of a GC SPME glass liner insert, however the added volume of the syringe was found to help with the condensation of water during sampling durations > 24-48 hours (depending on flow rate).

### Cell Culture

SK-OV-3 ovarian adenocarcinoma cells (human origin; HTB-77) were obtained from ATCC. Cells were maintained in RPMI 1640 (Sigma-Aldrich, St. Louis, MO, USA) supplemented with 10% FBS (ATCC) and 1% penicillin (100 I.U. / mL; ATCC) / streptomycin (100 µg/mL; ATCC). Cells were maintained in a humidified incubator at 37°C and 5% CO_2_. Cultures were grown in standard T-75 flasks and passaged at 70-85% confluency.

To seed the Biodome for VOC analysis, approximately 500,000 cells – estimated using a hematocytometer – were seeded in the Biodome. Cells were allowed to adhere from within a standard incubator (see above) for 24 hours, and the culture media was replaced immediately before the start of analysis. Vented caps from standard T-25 flasks (VWR) were allowed to rest on top of the inlet and outlet of the Biodome – maintaining an enclosed environment and CO_2_ exchange with the media. The total media volume in the Biodome was maintained at 5 mL for all experiments. To limit the background signal from the culture media, dialyzed fetal bovine serum (FBS) was used in place of traditional FBS, ensuring small molecule (<10,000 Da) concentration was reduced. Finally, to improve the consistency of heating and reduce condensation, the Biodome was completely submerged within a bead bath maintained at 37°C. Gas flow was set 24 hours before the start of experimentation to ensure an equilibrium was reached within the two-stage regulator.

### Bacterial Culture

DH5α *E. coli* were transformed with plasmid containing the ampicillin resistance gene (pUC19, Addgene Plasmid #50005) and streaked on LB agar plates spiked with 100 µg/mL ampicillin. Isolated colonies were inoculated in 100 µg/mL ampicillin spiked LB broth and grown for 24 hours. OD_600_ measurements were taken and used to dilute the bacterial culture such that the initial OD_600_ seeding concentration was approximately 0.1 (true value = 0.099) at the start of VOC collection. Bacteria were seeded into the Biodome containing a total of 5 mL of LB broth spiked with 100 µg/mL ampicillin and TDU tubes were replaced every 24 hours. The carrier gas composition was 95% air / 5% CO_2_, to allow compatibility with cellular analysis, and a flow rate of 18 mL/min was maintained for the duration of sampling. Gas flow was set 24 hours before the start of experimentation to ensure an equilibrium was reached within the two-stage regulator.

### Fluorescent Imaging

Prior to the start of the live/dead assay, seeded cells were allowed to adhere for 24 hours in a humidified incubator at 37°C and 5% CO_2_ and culture media was replaced immediately before introduction to the flow system. The Biodome was then incubated in the flow system for durations spanning 0 and 4 days. Following growth in the flow system, live/dead fluorescent probes (Blue/Green ReadyProbes, Invitrogen) were added to the Biodome according to the manufacturer guidelines. In brief, 2 drops of the ready-to-use stain were added per millimeter, totaling 8 drops. The biodome was then placed in a humidified incubator at 37°C and 5% CO_2_ for 30 minutes. After 30 minutes, the Biodome was imaged using Eclipse Ts2R microscope (Nikon, Minato City, Tokyo, Japan) with a Coolsnap Dyno CCD (Photometrics, Tucson, Arizona, USA) and 10x objective. A FITC filter (EX: 482 nm, EM: 536 nm; Nikon) was used to excite the live stain (Hoechst 33342) while a DAPI filter (EX: 390 nm, EM: 475 nm; Nikon) was used to visualize the dead stain. Fluorescent image processing and cell counting was performed using ImageJ.

To evaluate cell morphology, actin fluorescence staining was performed in live cells using CellMask™ Orange Actin Tracking Stain (Invitrogen). Cells were grown out in the specified incubation system for approximately 3 days until the confluency had reached 70-80%. 1000x stock solution of the actin tracking stain in DMSO was diluted to 1x in the RPMI1640 culture media described previously. Live/dead fluorescent probes were concurrently added to the staining media at a ratio of 2 drops per milliliter and cells were incubated for 30 minutes in a humidified incubator as previously described. Cells were rinsed twice with a wash buffer comprised of culture media previously equilibrated in a humidified incubator at 5% CO_2_ and 37°C for 30 min. Cells were imaged using a Leica DMi8 (Leica, Wetzlar, Germany) microscope with DFC345 FX monochrome camera (Leica) and Texas Red (EX: 560 nm, EM: 645 nm), DAPI, and FITC filters to visualize the actin, dead cell nuclei, and live cell nuclei, respectively. Images were processed and fluorescent channels overlaid using ImageJ. All fluorescent images were captured using cells at passage < 10 using a 10x objective.

### Stable isotope labeling of metabolites in cell culture

RPMI 1640 base medium without sodium bicarbonate and glucose was used (Sigma Aldrich), supplemented with 2 g / L ^13^C_6_–D-glucose (Cambridge Isotope Laboratories) and sodium bicarbonate (Sigma Aldrich). The final media contained 10% dialyzed FBS and 1% penicillin / streptomycin. To ensure equivalency when assessing isotopically labeled volatilomes, SK-OV-3 cells were passaged 20 times in their respective medias. Labeled and unlabeled cells were independently seeded into the glass culture vessel and allowed to grow out for 24 hours before starting sampling. Prior to connecting the Biodome to the flow system, culture media was replaced to ensure previous accumulation of VOCs did not influence results.

### 13C Peak Alignment

^13^C_6_-D-glucose was chosen as the labeling method because it allows peak alignment using retention times, given ^13^C and ^12^C isomers elute from the column at the same time^50, 51^. Chromatograms were first aligned within each experimental condition (i.e., labeled, and unlabeled) using ChromaTOF Statistical Compare package. Then, to assist with the identification of ^13^C-labeled VOCs, custom R code was developed (https://github.com/BSmithLab/Biodome) to facilitate alignment of unlabeled and labeled peaks by providing external control over the retention time variability. For each given peak in the unlabeled condition, all ^13^C-labeled peaks that fell within the allowed retention time window were identified, regardless of their underlying mass spectrum. This approach was deemed necessary to ensure chromatographic alignment between labeled and unlabeled conditions did not apply spectral deconvolution, which influences the resulting mass spectrum assigned to each peak. Mean retention times were determined for all ^13^C labeled and unlabeled peaks, and the first-dimension retention time threshold was set at 10 seconds while the second-dimension was set to 0.07 seconds. Labeled and unlabeled mass spectrums were then manually compared.

### RNA inhibition

Pooled siRNA oligonucleotides were purchased from Dharmacon (Lafayette, CO, USA) targeting *CPT2* [siGENOME SMARTpool Human CPT2; 5’ – GGCAGAAGCUGAUGAGUAG – 3’, 5’ – UGGCAUACCUGACCAGUGA – 3’, 5’ – GGAAAGUGGACUCGGCAGU – 3’, 5’ -CAAGAGACUCAUACGCUUU] and a non-silencing (sham) control [siGENOME Non-Targeting siRNA Pool #1; 5’ – UAGCGACUAAACACAUCAA – 3’, 5’ – UAAGGCUAUGAAGAGAUAC – 3’, 5’ – AUGUAUUGGCCUGUAUUAG – 3’, 5’ – AUGAACGUGAAUUGCUCAA – 3’]. SK-OV-3 cells were grown in a 6-well format and transfected at 80% confluency with complexes consisting of 50 nM pooled RNAi probes and 2 µg/mL Lipofectamine^TM^ 2000 (Invitrogen) in Opti-MEM^®^ I Reduced Serum Medium (ThermoFisher, Waltham, MA, USA). In the 6-well format, an extra well was allocated for direct seeding into the Biodome for VOC analysis. After 24 hours in a humidified incubator at 37°C and 5% CO_2_, the media was aspirated and the transfected cells were trypsinized. Approximately 500,000 cells were then seeded into the Biodome containing 5 mL of RPMI1640 with 10% dialyzed FBS and 1% Pen/Strep. The cells were allowed to adhere for roughly 1 hour in an incubator before being transferred to the flow system. For the transient RNAi assay, media was replaced after 24 hours, defining the 0-hour time point.

### RNA Extraction

Following RNAi, cellular RNA was extracted using PureLink^®^ RNA Mini Kit (Life Technologies, Carlsbad, CA, USA) according to manufacturer guidelines. In brief, fresh lysis buffer containing 1% 2-mercaptoethanol (Sigma Aldrich) was used to lyse the cells directly in the 6-well plate. Lysate was homogenized using 10 repeat passes through an 18-gauge needle (BD, Franklin Lakes, NJ). Binding, washing, and elution of purified RNA was performed according to kit specifications. Purity was assessed using absorbance values from a NanoDrop^TM^ One (ThermoFisher Scientific) and all samples were found to have A_260_/A_280_ ratios between 1.98 – 2.06. Samples were stored at –80°C.

### RT-qPCR

Purified RNA samples were thawed on ice and subject to reverse transcription using the QuantiTect Reverse Transcription Kit (Qiagen, Germantown, MD, USA). In brief, 100 ng of RNA was used as the starting weight. The genomic DNA elimination and reverse transcription reactions were prepared according to kit specifications. Reverse transcription reaction was allowed to proceed for a total of 15 minutes. Samples were stored for less than two weeks at –20°C before proceeding to qPCR.

The resulting cDNA was then subject to quantitative PCR to determine expression of *CPT2* and housekeeping gene *GAPDH*. Primers for *CPT2* [RefSeq Accession: NM_000098; forward primer 5’ – TGGTTTATCTGCCCGTATGC – 3’ and reverse primer 5’ – TGCTCCAAGTACCATGGC – 3’] and *GAPDH* [RefSeq Accession: NM_002046; forward primer 5’ – ACATCGCTCAGACACCATG – 3’ and reverse primer 5’ – TGTAGTTGAGGTCAATGAAGGG – 3’] were purchased from IDT (Newark, NJ, USA). The qPCR reaction was prepared in a 96-well format (ThermoFisher Scientific) using iTaq Universal SYBR Green Supermix (Bio-Rad Laboratories, Hercules, CA, USA), according to manufacturer guidelines. In brief, a 20 uL reaction size was selected, using 500 nM of the forward and reverse primer, 1 uL of cDNA, 10 uL of 2x iTaq, and remaining volume balanced with nuclease-free H_2_O. 96-well plates were transferred to a qTOWER 2.0 instrument and expression quantified using the following cycling conditions: 50°C for 2 minutes, 95°C for 5 minutes, followed by 40 cycles of amplification at 95°C for 15 seconds then 60°C for 1 minute. Annealing at 72°C was allowed to proceed for 10 minutes, at which point the dissociation characteristics were assessed by ramping the temperature from 65°C to 95°C using a step size of 2°C, equilibration time of 6 seconds, and heating rate of 0.5°C/second. Independent biological replicates were analyzed in technical triplicate and averaged.

### GC×GC-TOFMS

Thermal desorption tubes (Carbopack C, Carbopack B, and Carbosieve SIII, see Results) containing retained VOCs were analyzed using comprehensive GC×GC-TOFMS (Pegasus 4D^®^, LECO Corp. St. Joseph, MI), equipped with an autosampler (Multipurpose Sampler RoboticPro^®^, Gerstel Inc., Linthicum Heights, MD). TDU tubes were stored for a maximum of 8 days at 4°C to limit degradation^49^ and analyzed in order of collection. Volatiles were desorbed into the TDU 2 (Gerstel Inc.) inlet for a total of 4 minutes using variable temperature programming and splitless desorption mode. The TDU 2 inlet was initially held at 50°C for 30 seconds after which a ramp rate of 700°C/min was applied until the final temperature of 300°C was achieved. The inlet temperature was held at 300°C for 3 minutes prior to the start of chromatographic analysis. The transfer temperature was maintained at 310°C. The TDU 2 inlet was paired with a cooled injection system (CIS) to enhance analyte sensitivity. The CIS was initially maintained at – 100°C for the first 30 seconds of the run. The CIS temperature was then ramped at a rate of 12°C/sec until the final temperature of 275°C was achieved. The CIS was then maintained at 275°C for 3 minutes.

The instrument was fitted with a two-dimensional column set consisting of a Rxi-624Sil MS (60 m × 250 µm × 1.4 µm; Restek^®^, Bellefonte, PA) first dimension column and a Stabilwax (1 m × 250 µm × 0.5 µm, Restek) second dimension column joined together by a press-fit connection (length × internal diameter × film thickness, respectively). The first-dimension column was set at an initial temperature of 50°C and held for 2 minutes. The temperature was increased at a rate of 5°C/min until reaching the target temperature of 230°C, where a final hold time of 5 minutes was applied. The secondary oven was maintained at a +5°C offset relative to the primary oven. A quad-jet modulator was used with a 2 second modulation period and a +15°C offset relative to the secondary oven. The transfer line was maintained at 250°C and the ion source at 200°C for the duration of the run. Helium (UHP, 99.999%) was used as the carrier gas at a flow rate of 2 mL/min. Mass spectra were acquired at 100 Hz over a mass range of 35-300 Da with an ionization energy of -70eV. Prior to all sample analysis, a PFTBA standard was run to tune the mass spectrometer. Empty vials (blanks) were run at the start of each sample set to monitor the system for contamination. An alkane standard (C8-C20; Sigma Aldrich, St. Louis, MO) was sampled at the conclusion of testing for use in determination of retention indices.

### Data Processing

Within individual sample chromatograms, subpeaks in the second dimension were required to meet a signal-to-noise ratio ≥ 6 to be combined. The signal-to-noise cutoff for peak selection was set at 20:1 for a minimum of two apexing masses. The baseline signal was drawn through the middle of the noise. To align peaks across chromatograms, the Statistical Compare software package in ChromaTOF^®^ Version 4.72.0.0 (LECO Corp.) was utilized. Peaks were allowed to shift by a maximum of 18 seconds (9 modulation periods) in the first dimension and 0.15 seconds in the second dimension. The resulting aligned peaks were compared to the National Institute of Standards and Technology (NIST) 2011 Mass Spectral Library and tentative peak names were assigned if the spectral similarity score was ≥ 600 (60%). A secondary round of peak picking was performed on aligned chromatograms using a signal-to-noise threshold of 5 and a minimum spectral similarity ≥ 60%. The resulting dataset was then manually filtered to remove poorly resolved peaks eluting before 400 seconds and known contaminants (siloxanes). Quantitative values for signal abundance were obtained by integrating peaks areas using the unique ion mass.

VOC identities were assigned according to the metabolomic reporting standards established previously^48^. Compounds received an ID confidence level between 1 and 4, with 1 representing the highest confidence supported by two independent and orthogonal data sources^48^. In brief, compounds with mass spectral match ≥ 85%, to the NIST 2011 Mass Spectral Library, received an initial confidence level of 3. Compounds below this mass spectral threshold were assigned an ID level of 4 and labeled as unknown. The tentative VOC name and functional group is reported for all compounds with an ID level ≤ 3. A C8-C20 alkane standard was analyzed (**Fig. S3**) and used to assign retention indices for all VOCs. An ID level of 2 was the highest classification in this study, verified with a ≥ 85% mass spectral match and a retention index that is consistent with the mid-polar Rxi-624Sil stationary phase using the mean of published RIs, according to an approach previously established^109, 110^. If mass spectral and polar second dimension chromatographic information supported the assigned of a functional group to a compound with an ID level of 4, then the compound was named by the functional group.

### Statistical Analysis

To focus on the reproducible aspects of the biological volatilomes, compounds variably present (> 20% missing observations) in the experimental and control volatilomes were removed, unless otherwise indicated. Integrated peaks were log_10_ transformed in R, Version 4.0.3 (The R Foundation for Statistical Computing, Vienna, Austria) and missing values were imputed to 0 abundance. Pairwise, two-sided students’ t-tests were applied comparing biological and control VOC abundances across the sampling duration. Abundance observations were assumed to be unpaired, with unequal variance. The resulting *p*-values were then adjusted according to the Benjamini-Hochberg procedure^53^ using the “p.adjust” function in the base R “stats” library.

### Sterilization and re-use Procedure

Following the conclusion of sampling and analysis in the Biodome, the TDT adapter was removed and submerged in a 100% ethanol bath for at least 1 hour and subsequently sonicated for 2 minutes. Removal of the live culture is dependent on the organism considered. For microbiological analysis, a freshly prepared 10% bleach solution was added directly through the outlet of the Biodome and left at room temperature for 15 minutes. For adherent cell culture, spent media was aspirated and 5 mL of 0.25% Trypsin / 0.53 mM EDTA was added to the Biodome and incubated at 37°C for 5-10 minutes to dissociate cells from the surface. Trypsinization was deemed a highly beneficial step to prevent accumulation of leftover cellular debris on the culture surface following autoclaving (later step). In both cases, the liquid culture was then emptied, and the Biodome was subject to two additional wash steps using bleach with significant lateral motion. Afterwards, the Biodome was flushed thoroughly with DI water and then autoclaved for 1 hour. Following sterilization in the autoclave, the biodome was then placed in an acid bath (3M HCl) overnight (≥ 10 hours), although 3 hours was found to be sufficient. Following the acid bath, the Biodome was flushed with DI water and transferred to a base bath (approximately 150 mM NaOH) for at least 3 hours. Finally, to reduce noise from the culture vessel, the Biodome was flushed again with DI water, placed in a glass beaker, and heat treated for > 12 hours at 100°C. The authors recommend treating the Biodome in the base bath second, to limit the detection of acid-related contaminants.

## Acknowledgements

The authors would like to acknowledge Drs. Heather Bean and Trenton Davis in the School of Life Sciences at Arizona State University for their project input and initial discussions as to design feasibility. The authors would like to give a special thanks to Christine Roeger for her glassblowing expertise and fabrication of the Biodome culture vessels utilized in this study.

## Data Availability

All raw chromatographic and mass spectral data have been made publicly available in the MetaboLights^111^ repository at: https://www.ebi.ac.uk/metabolights/MTBLS8252

## Code Availability

Custom R code for the alignment of ^12^C and ^13^C-labeled VOCs and for the statistical analysis of SK-OV-3 and *E. coli* volatilomes has been made publicly available in the Smith Lab Github repository (https://github.com/BSmithLab/Biodome).

