## Supplementary Information for "An engineered culture vessel and flow system to improve the *in vitro* analysis of volatile organic compounds"

**This PDF file includes:**

Figs. S1 to S8
Table S1

**Other Supplementary Materials for this manuscript include the following:**

Supplemental Movies 1 and 2
Supplementary Dataset 1 to 4

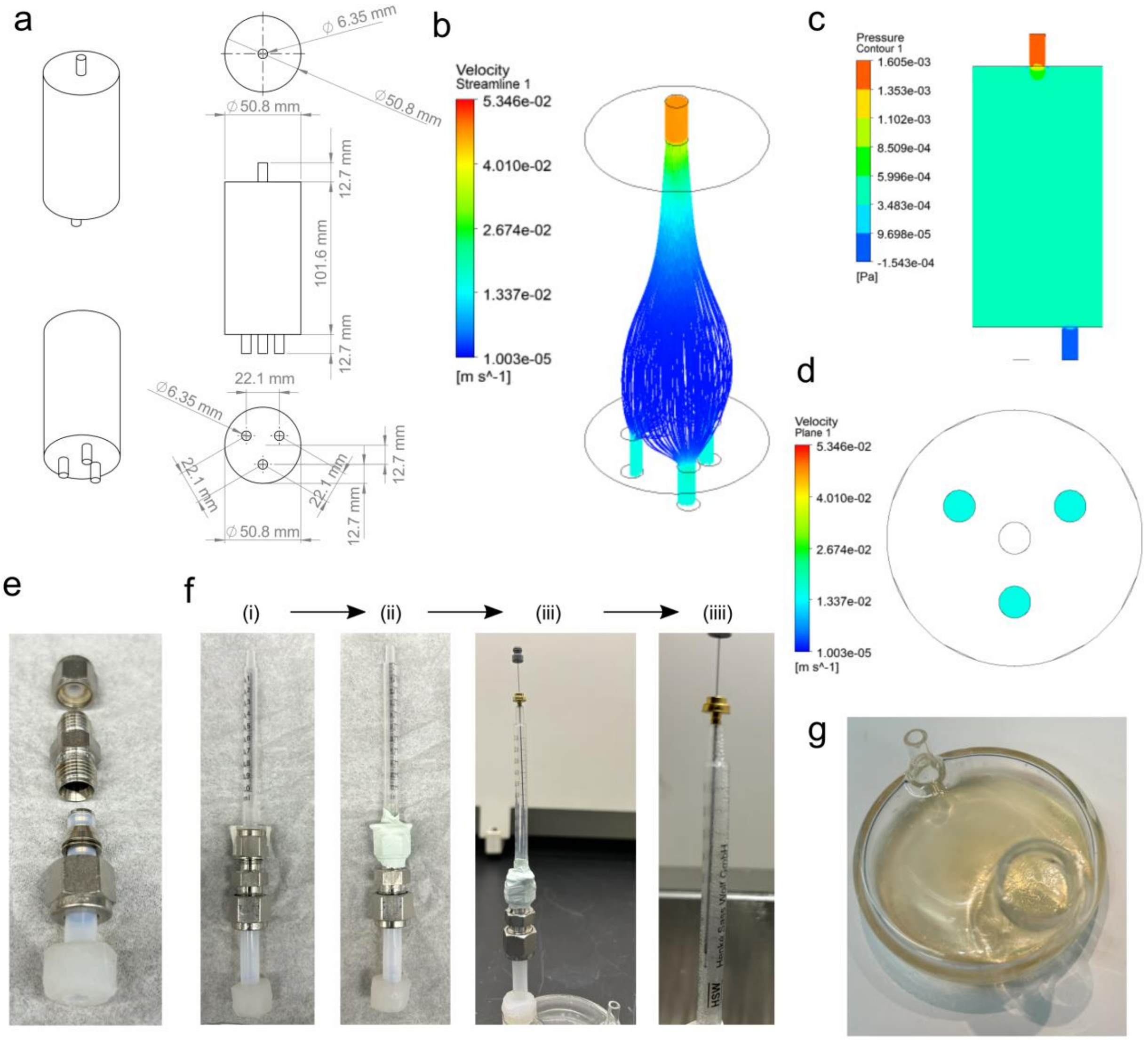


**Fig. S1. Biodome variations and adaptations.** (a) Schematic presentation of a multiplexed part designed in SOLIDWORKS® 2019 to allow experimental multiplexing, enhancing throughput. Fluid modeling was performed in ANSYS Fluent® 2019 R3 and (b) streamline plots, (c) pressure contours, and (d) velocity contours demonstrate desirable and equivalent flow rates at each of the outlets. (e) Individual components comprising the inexpensive TDT adapter. (f) Image series showing construction of the SPME adapter variant with: (i) minimal adhesive tape used to secure the syringe body in line with the flow path, (ii) use of oxygen-compatible PTFE tape (Restek) to seal the interface, (iii) insertion of the assembly into the Biodome device with the SPME allowed to rest naturally at the top of the syringe with sorbent material exposed, (iiii) visible condensation should form along the added gas flow path, with the image showing condensation following a 48 hour sampling duration. (g) Further adaptation to consider growth of ampicillin resistant *E. coli* on 6 mL solid LB agar containing 100 ug/mL ampicillin inside the Biodome. 50 uL of glycerol stock ($OD_{600}$ ~ 0.6) was added directly to the cooled agar and streaked using glass plating beads (Zymo Research, Irvine, CA, USA).

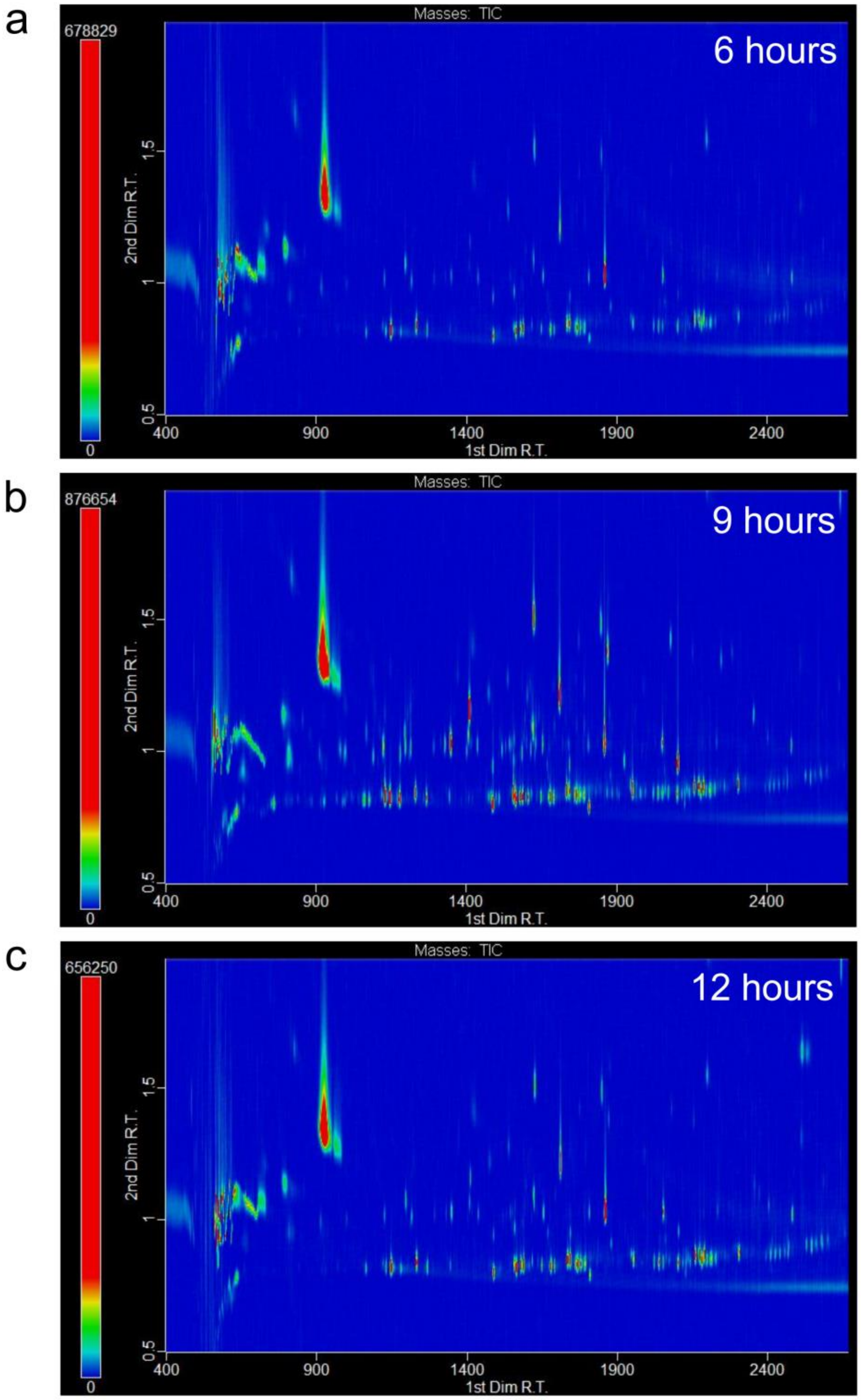


**Fig. S2. Variable sampling duration raw chromatograms.** Sampling duration < 24 hours were also considered, with representative chromatograms shown for (a) 6 hours, (b) 9 hours, and (c) 12 hours.

a

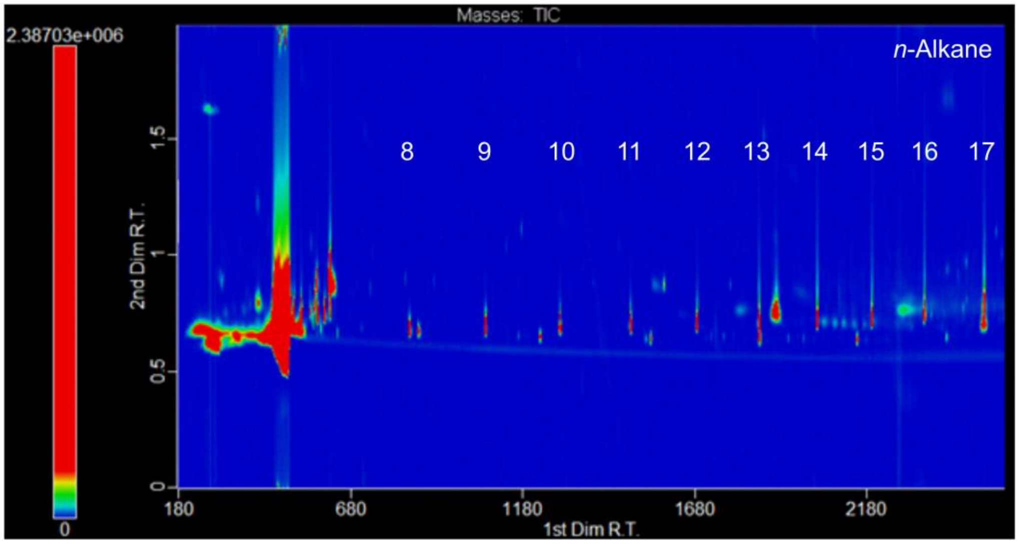


b

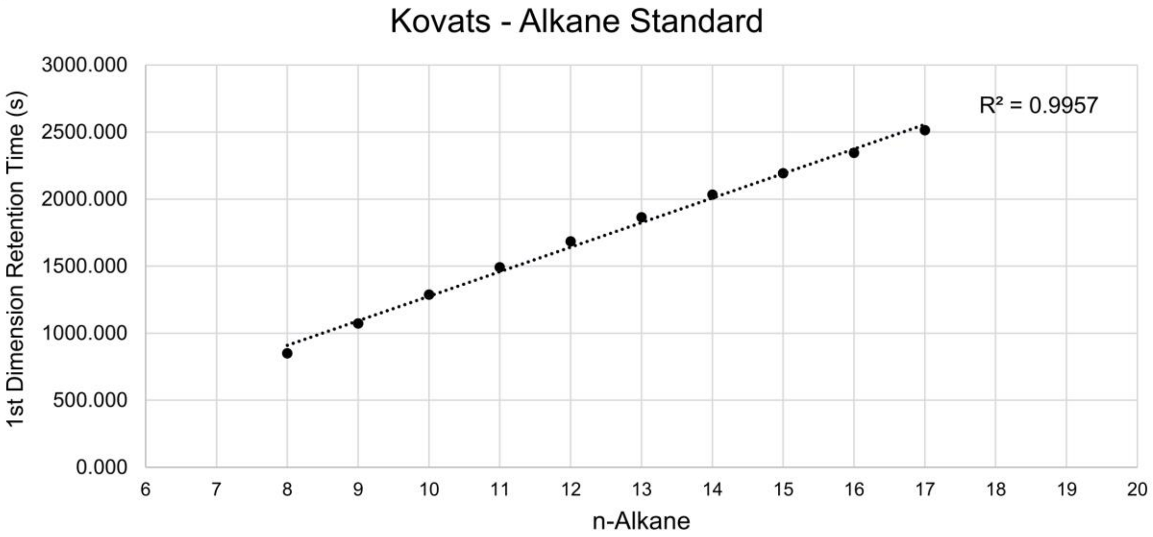


**Fig. S3. Alkane standard for determination of retention indices.** (a) Raw chromatogram from a C8-C20 alkane standard (Sigma Aldrich) showing octane to heptadecane. (b) Elution of alkanes shows a high degree of linearity in the first dimension, allowing extrapolation of retention indices that fell outside the range of the alkane standard.

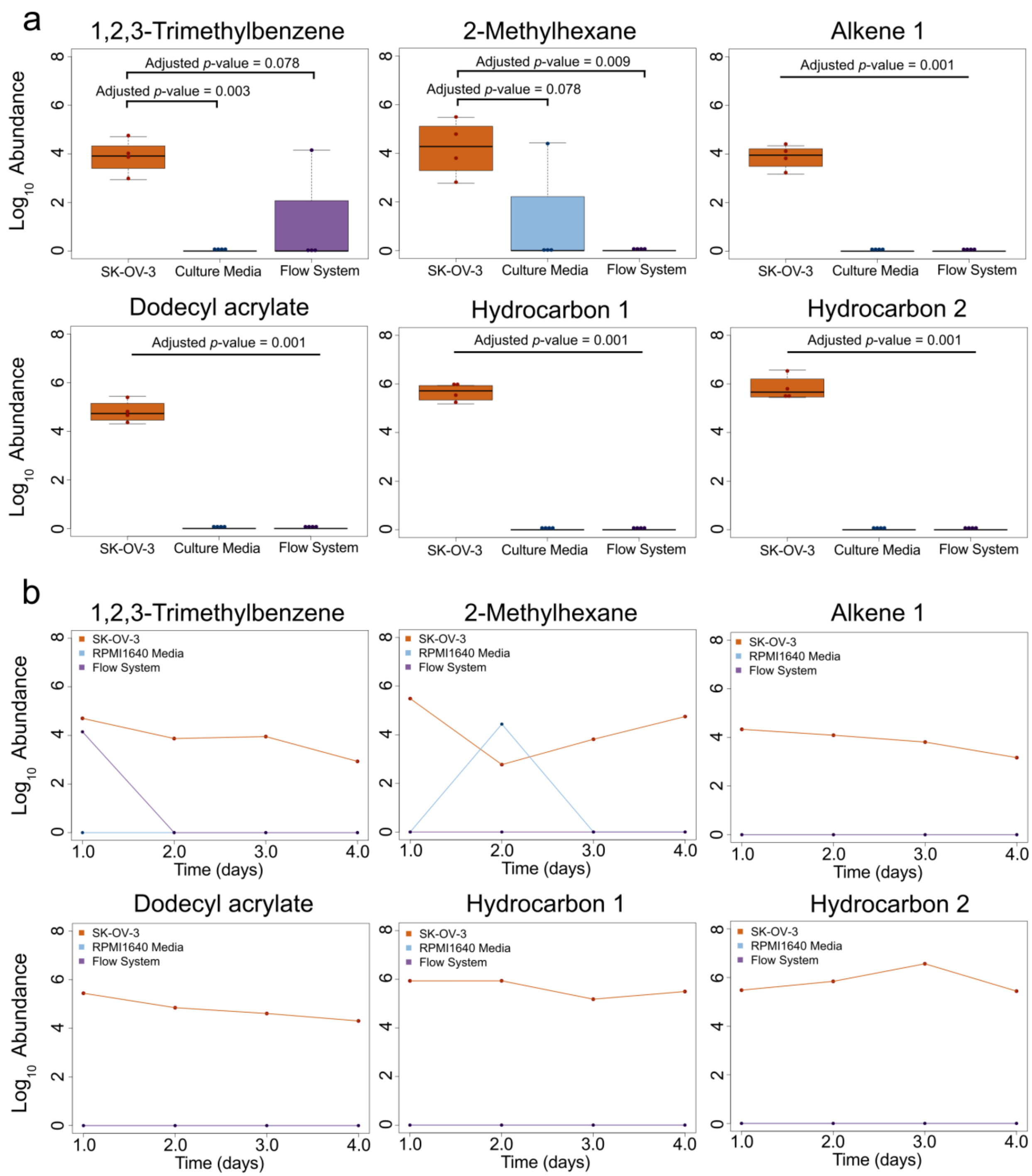


**Fig. S4. Additional VOCs uniquely observed in the SK-OV-3 volatilome.** Six additional VOCs presented in Table 1 are shown as (a) boxplots and (b) line plots. Statistically significant differences in the transformed peak abundances were observed for all higher mass VOCs (Alkene 1, Dodecyl acrylate, Hydrocarbon 1 and 2), indicating endogenous origin. Lower mass VOCs 1,2,3-trimethylbenzene and 2-Methylhexane were observed to have a single spurious aligned peak from both controls, suggesting endogenous origin, but may be related to typical resolution limitations at the beginning of the chromatographic run.

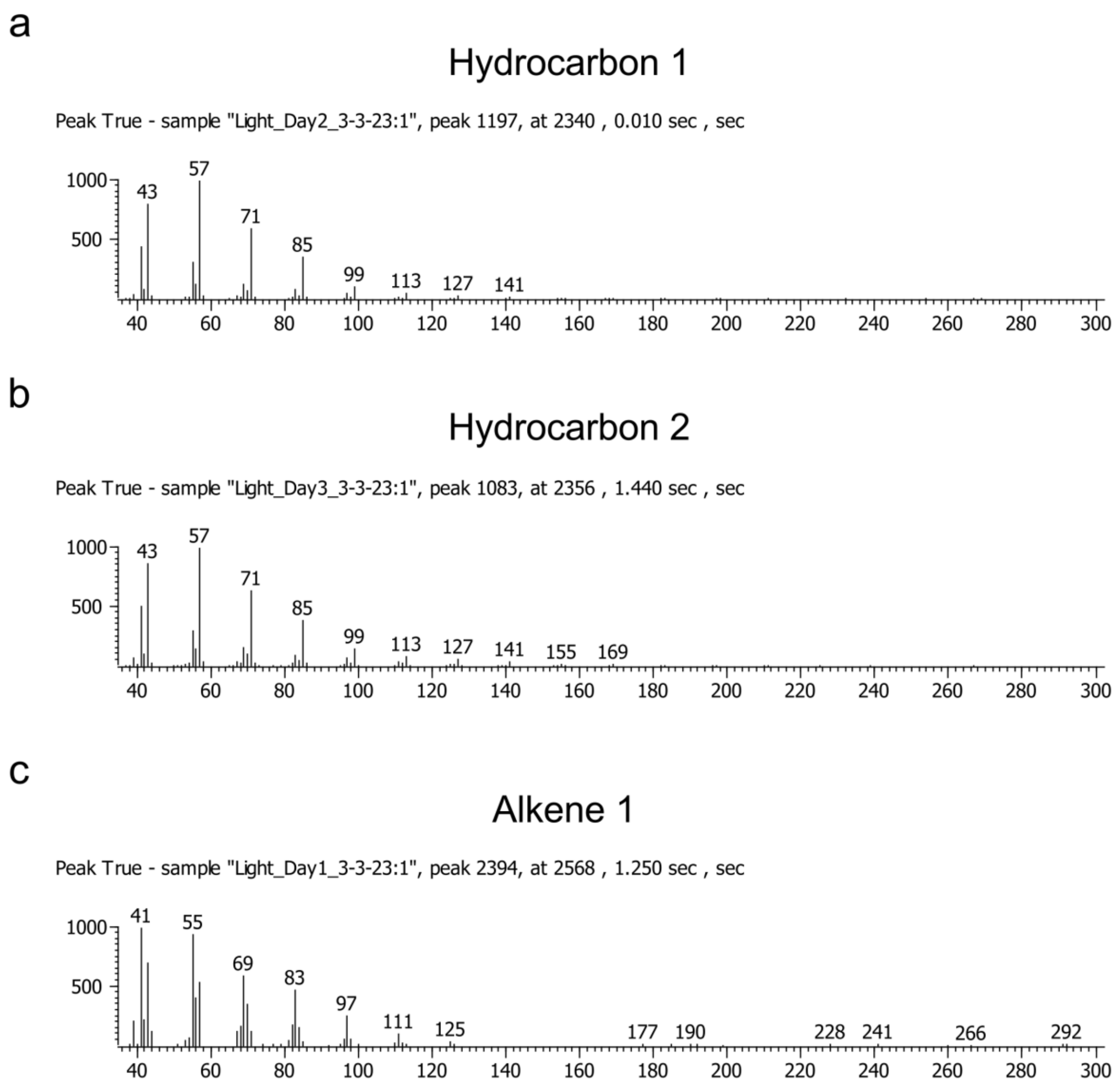


**Fig. S5. Supplemental mass spectrums supporting identification as two alkanes and an alkene.** Mass spectrums were selected using the aligned peak with the highest signal-to-noise ratio for (a) Hydrocarbon 1 eluting at 2336 seconds (average $^1t_R$), (b) Hydrocarbon 2 eluting at 2355 seconds (average $^1t_R$), and (c) Alkene 1 eluting at 2569 seconds (average $^1t_R$).

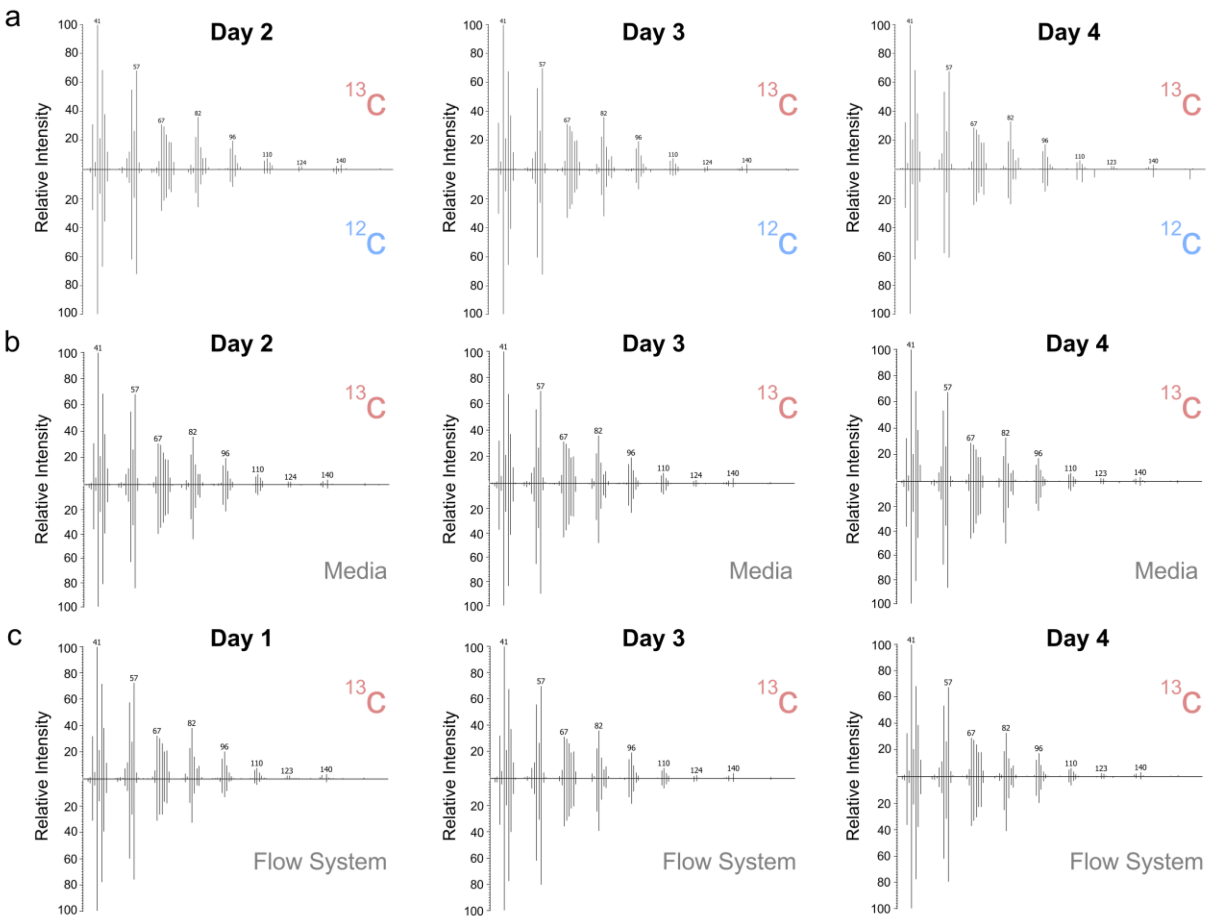


**Fig. S6. Supporting mass spectra comparing $^{13}$C-labeled and unlabeled 2-Decen-1-ol in controls.** 2-Decen-1-ol was observed across all days of sampling, including media and flow system controls. Mass spectra are paired by day of sampling comparing (a) $^{13}$C-labeled and unlabeled SK-OV-3 cells, (b) labeled cells and media control, and (c) labeled cells and flow system control. Fragments of *m/z* 59, 103, 107, 114, 125, 126, 127, 133, 139, 142, 143, and 147 were observed in 1 or fewer control spectra – indicating SILMC methodology in the Biodome has the ability to validate endogenous origin even when initially present in the exogenous volatilome.

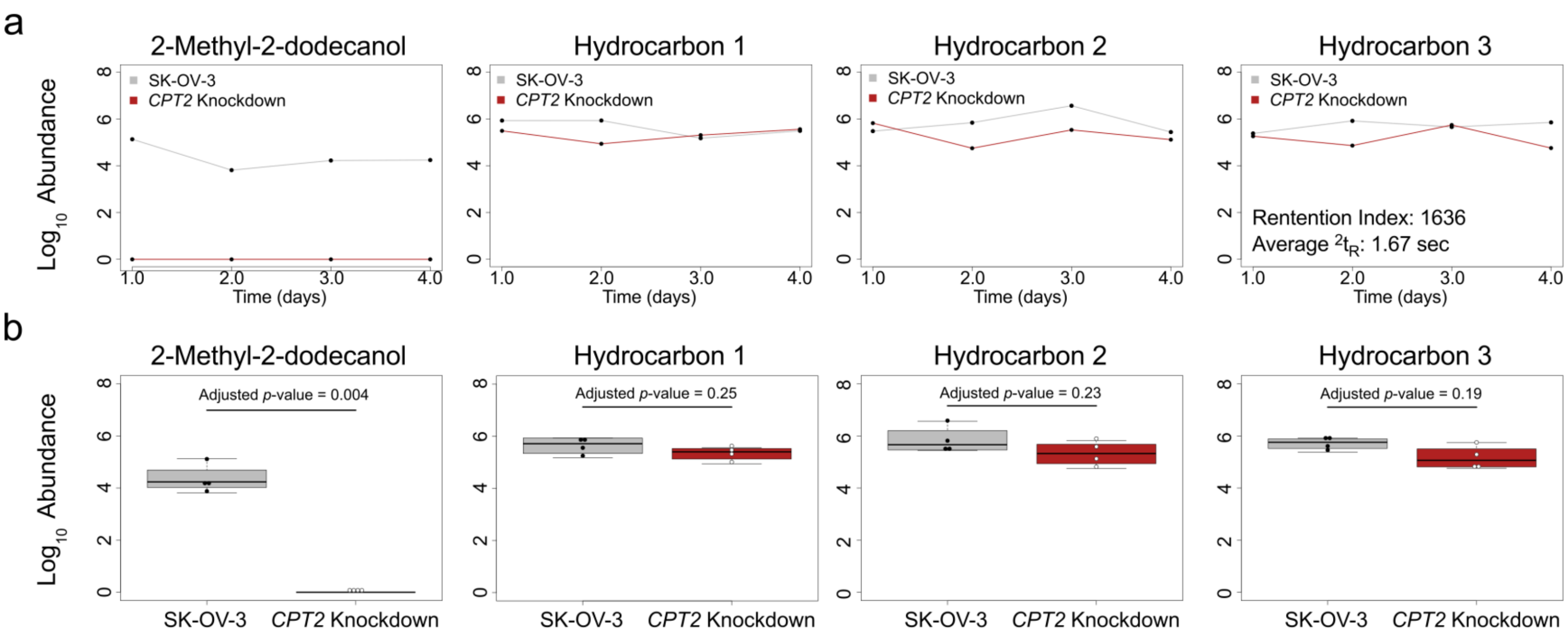


**Fig. S7. Plots supporting endogenous production of VOCs derived from fatty acid metabolism.** Four additional VOCs downregulated following RNA inhibition of the beta-oxidation pathway as (a) line plots and (b) boxplots. Hydrocarbon 3 was not initially included in the SK-OV-3 specific volatilome as one spurious aligned peak was identified in both the media and system controls.

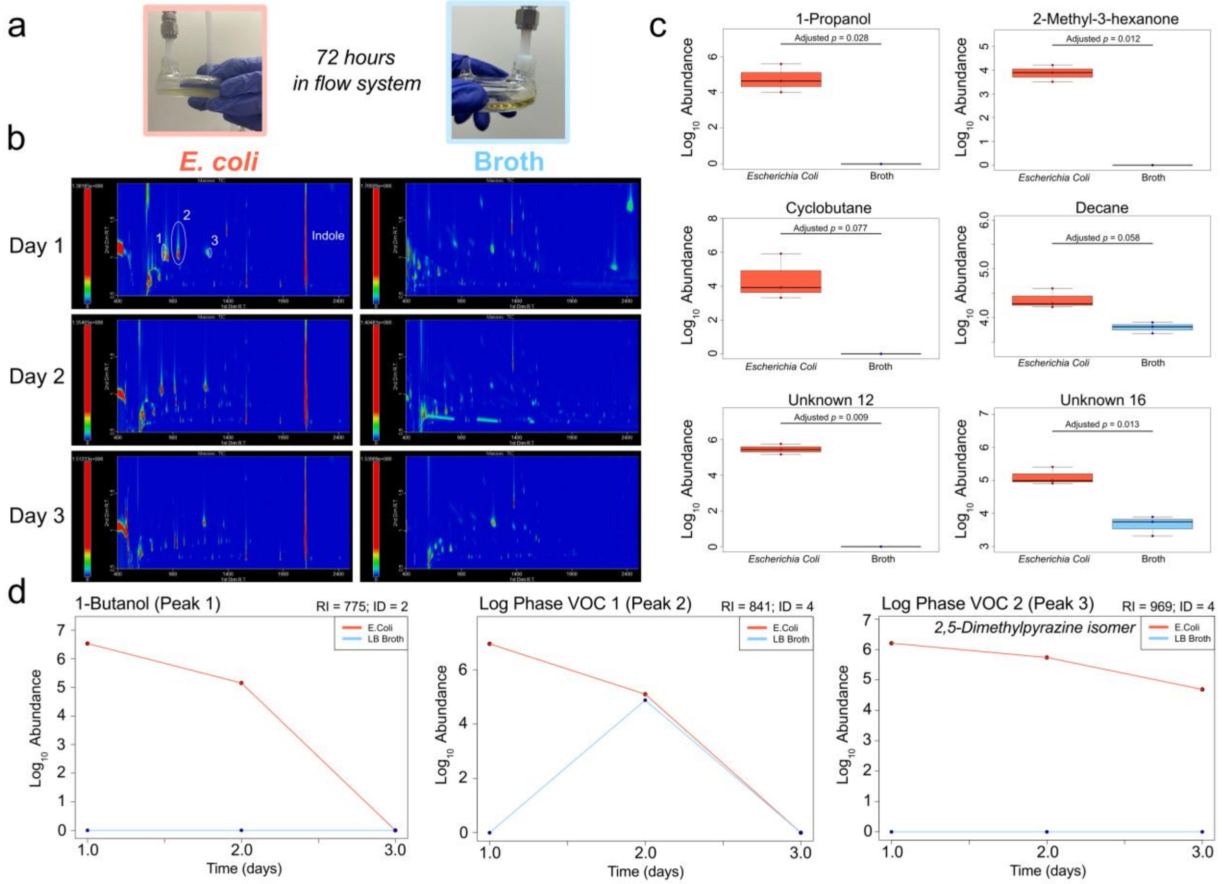


**Fig. S8. *Escherichia Coli* raw chromatograms and endogenous VOCs.** (a) Images showing (*left*) growth of *E. Coli* in the Biodome tool and (*right*) LB Broth control after 72 hours of sampling in the flow system. (b) Raw GC×GC-TOFMS chromatograms showing the reproducible nature of the Biodome tool for microbiological analysis. Indole abundance was estimated by summing integration slices in the first-dimension retention time window of 2092-2110 seconds. (c) Six representative VOCs with integrated peak abundances supporting endogenous production in *Escherichia Coli*. (d) Line plots showing VOCs elevated during the log phase of growth (0-24 hours). Retention index and Metabolomic Standards Initiative[48] ID confidence values are provided.

**Table S1. Full *Escherichia Coli* volatilome.** The compound name is presented first, followed by the functional group class. The *p*-value was calculated using a two-tailed student's t-test, comparing the LB broth and *Escherichia coli* volatilomes post-filtering and adjusted using the FDR procedure in R. The first ($^1t_R$) and second ($^2t_R$) dimension retention times are included, with the retention index calculated using a C8-C20 alkane standard mix. The VOC naming confidence level is reported according to previously established standards and VOCs uniquely detected in the broth or *E. Coli* conditions are indicated.

| VOC name | Chemical Class | *p*-value | FDR adjusted *p*-value | Avg 1tR (s) | Avg 2tR (s) | RI | ID Level | Specific to *E. Coli*? | Specific to LB Broth? |
|---|---|---|---|---|---|---|---|---|---|
| Acetone | Ketone | 0.48391 | 0.59389 | 639 | 0.67 | 652* | 2 | | |
| Methacrolein | Aldehyde | 0.01554 | 0.04196 | 702 | 0.69 | 687* | 2 | Yes | |
| 1-Propanol | Alcohol | 0.00920 | 0.02777 | 709 | 1.02 | 691* | 2 | Yes | |
| Unknown 1 | | 0.16261 | 0.33135 | 726 | 1.02 | 699* | 4 | | |
| Cyclobutane | Hydrocarbon | 0.03062 | 0.07690 | 799 | 1.06 | 758* | 3 | Yes | |
| Unknown 2 | | 0.83553 | 0.86766 | 804 | 0.77 | 763* | 4 | | |
| Unknown 3 | | 0.00600 | 0.01905 | 851 | 0.77 | 801 | 4 | Yes | |
| 2,3,5-Trimethylhexane | Hydrocarbon | 0.52371 | 0.62846 | 994 | 0.65 | 865 | 3 | | |
| Heteroaromatic 1 | Heteroaromatic | 0.32603 | 0.51782 | 1000 | 1.19 | 868 | 4 | | |
| 2,3-Hexanedione | Ketone | 0.23262 | 0.43700 | 1003 | 0.85 | 869 | 2 | | |
| 2,3,4-Trimethylhexane | Hydrocarbon | 0.58792 | 0.66837 | 1028 | 0.66 | 880 | 3 | | |
| Nonane | Hydrocarbon | 0.52000 | 0.62846 | 1084 | 0.65 | 906 | 2 | | |
| Unknown 4 | | 0.00315 | 0.01308 | 1111 | 0.93 | 918 | 4 | Yes | |
| Hydrocarbon 1 | Hydrocarbon | 0.34124 | 0.52423 | 1123 | 0.66 | 924 | 4 | | |
| 5-Methyl-2-hexanone | Ketone | 0.59734 | 0.67201 | 1141 | 0.88 | 932 | 2 | | |
| Hydrocarbon 2 | Hydrocarbon | 0.35899 | 0.52423 | 1153 | 0.66 | 937 | 4 | | |
| 2-Methyl-3-hexanone | Ketone | 0.00273 | 0.01178 | 1156 | 0.82 | 939 | 3 | Yes | |
| Unknown 5 | | 0.93452 | 0.94326 | 1172 | 0.84 | 946 | 4 | | |
| Unknown 6 | | 0.00080 | 0.00942 | 1173 | 0.85 | 947 | 4 | Yes | |
| Unknown 7 | | 0.00526 | 0.01834 | 1189 | 0.80 | 954 | 4 | Yes | |
| 2,5-Dimethylpyrazine | Heteroaromatic | 0.55969 | 0.64996 | 1199 | 1.11 | 959 | 2 | | |
| Hydrocarbon 3 | Hydrocarbon | 0.54027 | 0.64120 | 1200 | 0.66 | 959 | 4 | | |
| Cyclohexanone | Ketone | 0.81501 | 0.85604 | 1232 | 1.05 | 974 | 2 | | |
| Hydrocarbon 4 | Hydrocarbon | 0.35919 | 0.52423 | 1257 | 0.66 | 986 | 4 | | |
| Unknown 8 | | 0.22779 | 0.43700 | 1257 | 0.94 | 986 | 4 | | |
| Unknown 9 | | 0.56788 | 0.65245 | 1265 | 0.67 | 989 | 4 | | |
| Hydrocarbon 5 | Hydrocarbon | 0.54649 | 0.64154 | 1273 | 0.66 | 993 | 4 | | |
| Unknown 10 | | 0.14855 | 0.31457 | 1285 | 0.86 | 999 | 4 | | |

| | | | | | | | | | |
|---|---|---|---|---|---|---|---|---|---|
| Aromatic Hydrocarbon 1 | Aromatic Hydrocarbon | 0.44057 | 0.56597 | 1303 | 0.87 | 1008 | 4 | | |
| Unknown 11 | | 0.36591 | 0.52692 | 1306 | 0.85 | 1009 | 4 | | |
| Decane | Hydrocarbon | 0.02216 | 0.05838 | 1326 | 0.67 | 1019 | 2 | | |
| Unknown 12 | | 0.00097 | 0.00942 | 1330 | 1.09 | 1021 | 4 | Yes | |
| Unknown 13 | | 0.23468 | 0.43700 | 1339 | 0.90 | 1025 | 4 | | |
| Unknown 14 | | 0.00926 | 0.02777 | 1346 | 0.86 | 1028 | 4 | | Yes |
| Dimethyl trisulfide | Other | 0.13505 | 0.29171 | 1358 | 1.09 | 1034 | 2 | | |
| Hydrocarbon 6 | Hydrocarbon | 0.35787 | 0.52423 | 1363 | 0.66 | 1037 | 4 | | |
| 2-Ethyl-6-methyl-pyrazine | Heteroaromatic | 0.00223 | 0.01047 | 1365 | 1.03 | 1038 | 2 | | Yes |
| Trimethylpyrazine | Heteroaromatic | 0.09405 | 0.21612 | 1369 | 1.06 | 1040 | 2 | | |
| 1,2,3-Trimethylbenzene | Aromatic Hydrocarbon | 0.79721 | 0.85604 | 1370 | 0.90 | 1040 | 2 | | |
| 2-Ethyl-5-methylpyrazine | Heteroaromatic | 0.39873 | 0.54631 | 1377 | 1.03 | 1044 | 2 | | |
| Unknown 15 | | 0.00417 | 0.01499 | 1381 | 0.92 | 1045 | 4 | | Yes |
| 2,2,7,7-Tetramethyloctane | Hydrocarbon | 0.45068 | 0.56597 | 1392 | 0.66 | 1051 | 3 | | |
| 2,2,4,4-Tetramethyloctane | Hydrocarbon | 0.72611 | 0.79212 | 1395 | 0.67 | 1053 | 3 | | |
| Hydrocarbon 7 | Hydrocarbon | 0.29302 | 0.51782 | 1403 | 0.66 | 1057 | 4 | | |
| Unknown 16 | | 0.00345 | 0.01330 | 1410 | 0.87 | 1060 | 4 | | |
| Hydrocarbon 8 | Hydrocarbon | 0.44618 | 0.56597 | 1412 | 0.71 | 1061 | 4 | | |
| Unknown 17 | | 0.38955 | 0.54631 | 1418 | 0.70 | 1064 | 4 | | |
| Heteroaromatic 2 | Heteroaromatic | 0.02280 | 0.05863 | 1423 | 1.17 | 1066 | 4 | | |
| Hydrocarbon 9 | Hydrocarbon | 0.71044 | 0.78294 | 1426 | 0.67 | 1068 | 4 | | |
| Aromatic Hydrocarbon 2 | Aromatic Hydrocarbon | 0.87950 | 0.90463 | 1435 | 0.93 | 1072 | 4 | | |
| 2,3,5,8-Tetramethyldecane | Hydrocarbon | 0.68969 | 0.76790 | 1438 | 0.67 | 1073 | 3 | | |
| Unknown 18 | | 0.00014 | 0.00884 | 1440 | 1.05 | 1075 | 4 | | Yes |
| Hydrocarbon 10 | Hydrocarbon | 0.35248 | 0.52423 | 1449 | 0.67 | 1079 | 4 | | |
| 3-Thiophenecarboxaldehyde | Other | 0.00118 | 0.00942 | 1451 | 1.73 | 1080 | 2 | | Yes |
| Unknown 19 | | 0.00030 | 0.00884 | 1451 | 0.99 | 1080 | 4 | | Yes |
| 2-Acetylthiazole | Azole | 0.05969 | 0.14014 | 1462 | 1.53 | 1085 | 2 | | |
| Unknown 20 | | 0.00058 | 0.00888 | 1473 | 1.05 | 1091 | 4 | | Yes |
| Unknown 21 | | 0.12935 | 0.28511 | 1477 | 0.69 | 1092 | 4 | | |
| 2-Methyl-3-isopropylpyrazine | Heteroaromatic | 0.43135 | 0.56128 | 1482 | 0.97 | 1095 | 2 | | |
| Benzyl alcohol | Functionalized Benzene | 0.00342 | 0.01330 | 1510 | 0.35 | 1109 | 2 | Yes | |

| | | | | | | | | | |
|---|---|---|---|---|---|---|---|---|---|
| Hydrocarbon 11 | Hydrocarbon | 0.31022 | 0.51782 | 1515 | 0.72 | 1112 | 4 | | |
| 3-Ethyl-2,5-dimethylpyrazine | Heteroaromatic | 0.17781 | 0.35561 | 1526 | 0.99 | 1118 | 2 | | |
| Unknown 22 | | 0.00561 | 0.01834 | 1529 | 1.19 | 1119 | 4 | | Yes |
| Hydrocarbon 12 | Hydrocarbon | 0.42411 | 0.55858 | 1534 | 0.68 | 1122 | 4 | | |
| 2-Hydroxybenzaldehyde | Other | 0.41592 | 0.55858 | 1534 | 1.54 | 1122 | 2 | | |
| Unknown 23 | | 0.31967 | 0.51782 | 1535 | 0.75 | 1122 | 4 | | |
| 1-Octanol | Alcohol | 0.81641 | 0.85604 | 1536 | 1.11 | 1123 | 2 | | |
| Heteroaromatic 3 | Heteroaromatic | 0.11925 | 0.26830 | 1539 | 0.99 | 1124 | 4 | | |
| Unknown 24 | | 0.00153 | 0.00969 | 1547 | 0.89 | 1129 | 4 | Yes | |
| Unknown 25 | | 0.39961 | 0.54631 | 1573 | 0.73 | 1142 | 4 | | |
| 2-Nonanone | Ketone | 0.00131 | 0.00942 | 1581 | 0.86 | 1146 | 2 | Yes | |
| Unknown 26 | | 0.00143 | 0.00962 | 1582 | 1.25 | 1147 | 4 | | Yes |
| Benzoic acid, methyl ester | Functionalized Benzene | 0.92060 | 0.93797 | 1605 | 1.23 | 1159 | 2 | | |
| Hydrocarbon 13 | Hydrocarbon | 0.98927 | 0.98927 | 1608 | 0.75 | 1160 | 4 | | |
| Unknown 27 | | 0.00048 | 0.00888 | 1620 | 0.92 | 1167 | 4 | | Yes |
| 2-Methylundecane | Hydrocarbon | 0.29822 | 0.51782 | 1628 | 0.70 | 1171 | 2 | | |
| Hydrocarbon 14 | Hydrocarbon | 0.39090 | 0.54631 | 1640 | 0.69 | 1177 | 4 | | |
| Unknown 28 | | 0.29056 | 0.51782 | 1659 | 1.38 | 1187 | 4 | | |
| 3,5-Diethyl-2-methylpyrazine | Heteroaromatic | 0.22544 | 0.43700 | 1666 | 0.94 | 1191 | 2 | | |
| 3,5-Dimethyl-2-cyclohexen-1-one | Ketone | 0.00188 | 0.00969 | 1683 | 1.18 | 1200 | 3 | | Yes |
| Unknown 29 | | 0.00054 | 0.00888 | 1692 | 0.95 | 1204 | 4 | | Yes |
| 2-Nonenal, (E)- | Aldehyde | 0.34501 | 0.52423 | 1729 | 0.99 | 1225 | 2 | | |
| Unknown 30 | | 0.00382 | 0.01424 | 1737 | 0.82 | 1230 | 4 | | Yes |
| Heteroaromatic 4 | Heteroaromatic | 0.30667 | 0.51782 | 1752 | 0.91 | 1238 | 4 | | |
| 2-Phenylpropenal | Functionalized Benzene | 0.00124 | 0.00942 | 1752 | 1.42 | 1238 | 2 | | Yes |
| 1-Phenyl-1-propanone | Functionalized Benzene | 0.31603 | 0.51782 | 1760 | 1.24 | 1242 | 2 | | |
| Benzyl nitrile | Functionalized Benzene | 0.46025 | 0.57134 | 1766 | 1.76 | 1246 | 2 | | |
| Unknown 31 | | 0.04845 | 0.11628 | 1772 | 1.39 | 1249 | 4 | | |
| 3-Butyl-2,5-dimethylpyrazine | Heteroaromatic | 0.32194 | 0.51782 | 1848 | 0.94 | 1291 | 2 | | |
| Unknown 32 | | 0.00178 | 0.00969 | 1865 | 1.16 | 1300 | 4 | | Yes |
| 1-Phenyl-2-butanone | Functionalized Benzene | 0.00033 | 0.00884 | 1881 | 1.21 | 1310 | 2 | Yes | |

| | | | | | | | | | |
|---|---|---|---|---|---|---|---|---|---|
| Unknown 33 | | 0.00087 | 0.00942 | 1901 | 1.12 | 1322 | 4 | | Yes |
| Unknown 34 | | 0.01279 | 0.03542 | 1915 | 0.86 | 1331 | 4 | | Yes |
| Nonanoic acid | Ester | 0.00546 | 0.01834 | 1932 | 0.08 | 1340 | 2 | | Yes |
| α-Ethylidene-benzeneacetaldehyde | Functionalized Benzene | 0.00181 | 0.00969 | 1946 | 1.06 | 1349 | 2 | | Yes |
| Heteroaromatic 5 | Heteroaromatic | 0.28494 | 0.51782 | 1950 | 0.92 | 1351 | 4 | | |
| Unknown 35 | | 0.00212 | 0.01041 | 1999 | 1.17 | 1380 | 4 | | Yes |
| 5-Butyldihydro-2(3H)-furanone | Ketone | 0.00973 | 0.02840 | 2001 | 1.30 | 1382 | 2 | | Yes |
| Unknown 36 | | 0.42201 | 0.55858 | 2004 | 0.93 | 1383 | 4 | | |
| Unknown 37 | | 0.00122 | 0.00942 | 2077 | 0.95 | 1428 | 4 | | Yes |
| Indole | Heteroaromatic | 0.00174 | 0.00969 | 2100 | 1.20 | 1442 | 2 | | |
| Ketone 1 | Ketone | 0.16239 | 0.33135 | 2112 | 0.85 | 1450 | 4 | | |
| 3-hydroxy-2,2,4-trimethylpentyl isobutyrate | Other | 0.04377 | 0.10744 | 2129 | 1.00 | 1460 | 3 | | |
| Unknown 38 | | 0.00130 | 0.00942 | 2240 | 0.90 | 1532 | 4 | Yes | |
| Unknown 39 | | 0.00030 | 0.00884 | 2261 | 0.31 | 1545 | 4 | Yes | |
| Unknown 40 | | 0.80965 | 0.85604 | 2267 | 0.70 | 1549 | 4 | | |
| Unknown 41 | | 0.00266 | 0.01178 | 2343 | 1.32 | 1599 | 4 | | Yes |
| Unknown 42 | | 0.01113 | 0.03163 | 2402 | 0.91 | 1634 | 4 | | Yes |

*Extrapolated RI from linear alkane series
